# Ambient light and mimicry as drivers of wing transparency in Lepidoptera

**DOI:** 10.1101/2020.06.26.172932

**Authors:** Mónica Arias, Jérôme Barbut, Rodolphe Rougerie, Manon Dutry, Mireia Kohler, Baptiste Laulan, Caroline Paillard, Serge Berthier, Christine Andraud, Marianne Elias, Doris Gomez

## Abstract

Transparency reduces prey detectability by predators. While transparent aquatic species hold higher transparency levels as the light availability of their habitat increases, less is known about such variation in terrestrial species. Lepidoptera species exhibiting transparent wings display various levels of transparency. Using two complementary approaches, we explore how the evolution of different transparency degrees relates to habitat openness, activity rhythm and mimicry syndrome (bee/wasp versus dead-leaf mimic). First, by exposing artificial moth-like prey to wild avian predators in a range of habitat openness, we show that survival is lower in more open habitats. We also found that less transparent morphs are more attacked than more transparent ones, regardless of habitat openness degree. Second, by analysing the evolution of wing features and ecological traits in 107 clearwing species, we found that diurnal species transmit more light than nocturnal species under certain conditions (when considering only forewings, at smaller clearwing surfaces and at larger wing lengths) and that species flying in open habitats and exhibiting large percentages of clearwing surface transmit slightly more light than those flying in closed habitats, although this is reversed at smaller percentages of clearwing surfaces. Additionally, bee/wasp mimics are more often diurnal and have higher and less variable light transmittances than dead-leaf mimics, which are more often nocturnal. Flying during the day, in open habitats and mimicking insects with transparent wings seem to promote high light transmittance under certain circumstances. Activity rhythm, habitat openness and species interactions play a crucial role in determining transparency design on land.

## Introduction

Transparency is common in aquatic environments where it reduces detectability by predators, especially in the pelagic environment where there is nowhere to hide (Johnsen 2014). The amount of light transmitted by transparent tissues varies between organisms. Experiments and modelling have shown that in brighter light conditions, the visual performance of aquatic predators is higher and only high levels of transparency confer protection to prey. Conversely, at dim light conditions that limit visual performance, a large range of transparency levels give equal protection (Figure S1, Johnsen and Widder 1998).

On land, transparency is present in only a handful of lineages, among which insects. In Lepidoptera, transparency is based on diverse microstructures (scales varying in morphology, insertion on the membrane, and colouration) and nanostructures (on scales and on the wing membrane), producing transmittance levels ranging from 10 to 90% (Gomez et al. 2021; Pinna et al. 2021). The triggers of the evolution of such diversity are still unexplored. Predation is likely important, as transparency efficiently reduces prey detectability (Arias et al. 2019, 2020; Mcclure et al. 2019). As in water, the visual performance of terrestrial predators is higher in bright than in dim light conditions, as shown in birds (Hodos et al. 1976; Kassarov 2003; Lind et al. 2013). We can thus hypothesize that diurnal terrestrial transparent species living in more open habitats - with brighter light conditions - would transmit more light compared to species flying at night and/or living in more closed habitats - with dimmer light conditions-. In addition, at low light availability, different levels of transmittance could confer similar protection against predators. Therefore, we expect that nocturnal and/or closed-habitat species would show larger variation in light transmittance, as neither high nor low transmittance levels are strongly selected.

Other visual anti-predatory defences might also be segregated according to light availability (activity rhythm and habitat openness). Predation risk can be reduced by mimicking unprofitable prey (batesian mimicry, Bates 1862) such as bees and wasps, or inedible elements such as leaves (masquerade, Skelhorn et al. 2010), species and items that are more common in habitats with different light conditions. Several small clearwing moths and butterflies exhibit yellow and dark stripes on their bodies and highly transparent wings (example in Fig. 1a), resembling harmful species for predators, such as bees and wasps, that often live in open habitats (Grundel et al. 2010; Yamaura et al. 2012), and bee/wasp mimics should do as well. Other clearwing Lepidoptera exhibit brownish colorations that combined with transparent surfaces that are presumably perceived as “holes” can resemble dead-leaves (example of a potential dead-leaf mimic in Fig. 1b) that are probably highly common in closed habitats were lots of dead leaves fall down. High light transmittance should be selected in bees, wasps and their mimics because the reduction of detectability already conferred by their small body sizes, can be enhanced by higher levels of transparency (Gomez et al. 2021), and because light conditions are brighter in open habitats. By contrast, dead-leaf mimics might show a larger variation in body size, and we expect that they would exhibit a broader diversity of light transmittance, as a range of transmittance levels in poor light conditions can contribute to mimic leaves in decomposition and confer similar protection against predators. The expected differences in light transmittance should also be associated to variation in morphology and coloration of wing scales, with absent or reduced scales for bee/wasp mimics, while all scale types should be probably found in dead-leaf mimics. Therefore, mimicry syndromes that are probably segregated in habitats with different light availability could also drive different transmittance optima in bee/wasp and in dead-leaf mimics.

**Figure 1.**
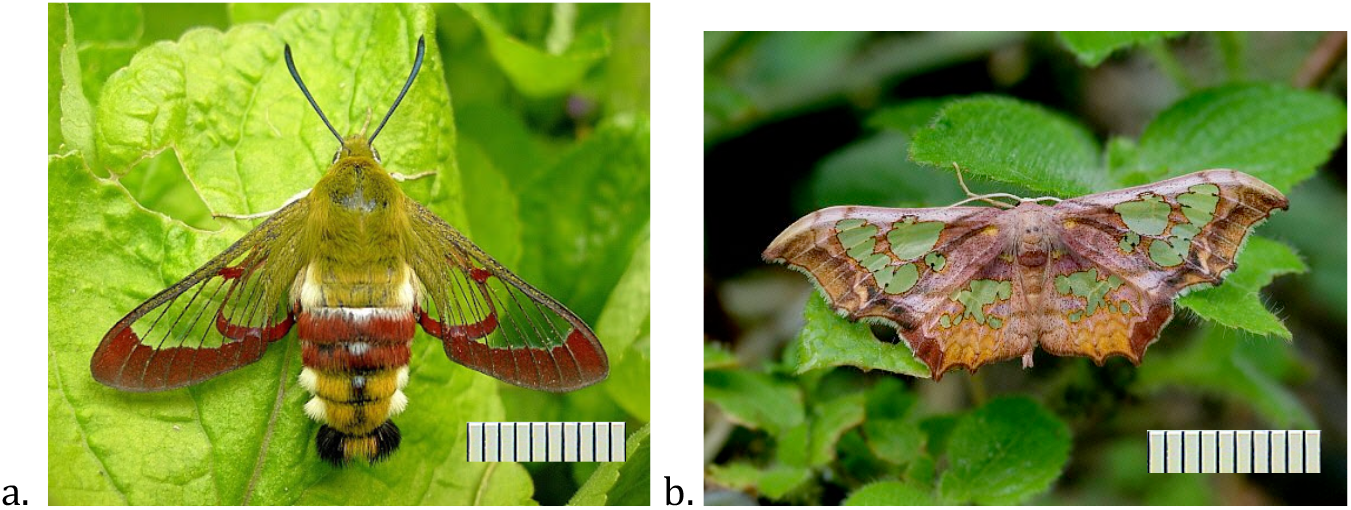
Examples of a bee/wasp mimic (a. adult female *Hemaris fuciformis fuciformis*, Catalonia, Spain. Photo: © Tony Pittaway) and a potential dead leaf mimic (b. *Pseudasellodes fenestraria*, Rio Kosnipata, Peru. Photo: © Adrian Hoskins). Forewing length is ~24mm in *Hemaris fuciformis fuciformis* and ~18mm in *Pseudasellodes fenestraria*. Each tick in the scale represents one mm.

Here, we investigate whether differences in transmittance found in Lepidoptera clearwing species are related to light availability and/or mimicry syndrome using two complementary approaches. First, we tested whether the difference in efficiency of different transmittance levels at reducing detectability is lower in closed habitat compared to open habitat, by carrying out a fieldwork experiment with artificial moth-like prey and natural predator communities. Second, using actual specimens of butterflies and moths, we tested whether differences in light transmission could be explained by species daytime rhythm, habitat and/or mimicry, by conducting comparative analyses at broad interspecific level. We took advantage of the large dataset of Lepidoptera species analysed by Gomez et al (2021), added species ecological traits (habitat, daytime rhythm and mimicry, known for 107 species) and explore the correlation and coevolution between them and light transmittance, and clearwing morphological (wing size, proportion of clearwing surface, and fore-/hindwing) and structural traits (scale characteristics). Together, these analyses help understanding the evolution of light transmission on terrestrial species exhibiting transparency.

## Material and Methods

### 1. Field experiment

### 1.a. Artificial butterfly Elaboration

To test for the efficiency of light transmittance at reducing detectability in habitats with different light availability, we elaborated plain grey artificial butterflies with paper wings and a malleable edible body (see Figure 2 for pictures of the artificial moths). Following the general methodology described in Arias *et al* (2020), butterflies did not replicate any real local butterfly, but mimicked a general grey moth with closed wings (i. e., a triangular shape of 25mm high by 36mm wide for a surface of 450mm^2^). Artificial moth colour was chosen based on the bark colour of evergreen oak (See ESM for further details). Artificial grey was chromatically similar, but lighter, to the evergreen oak trunk. This achromatic mismatch allowed us to test transparency as crypsis enhancer in imperfectly cryptic artificial prey. Moths were built combining mat laminated paper wings (as described in ESM). Four types of artificial butterflies were used, here listed in order of increasing proportion of light transmittance: completely opaque butterfly “C”; poorly transparent butterfly with 6 layers of transparent film “T6”; highly transparent butterfly with a single layer of transparent film “T1”; and fully transparent butterfly with no film in the transparent zones “T0”. For the butterflies that included transparent elements, two triangular windows of 252mm^2^ each (56% of total artificial moth surface) were cut down from the grey triangle and the remaining part was either left empty (T0) or glued on top of one (T1) or six (T6) layers of 3M of transparency film for inkjet printing. This transparent film was chosen as it is highly transparent, even in the UV range of the spectrum, and going from 1 to 6 transparent layers permitted a reduction of 50% of the transmittance between treatments (Fig. S2). The upper part of transparent areas was coated with a transparent matte varnish to reduce its shininess, making them more similar to real Lepidoptera transparent wings, which often exhibit nano antireflective structures (Pinna et al. 2021; Pomerantz et al. 2021). We added an edible malleable grey body on top of the wings that permitted registering and distinguishing bird, insect and mammal marks. Further details in the measurement of colour and transparency spectra, as well as in the fabrication of prey bodies can be found in ESM.

**Figure 2.**
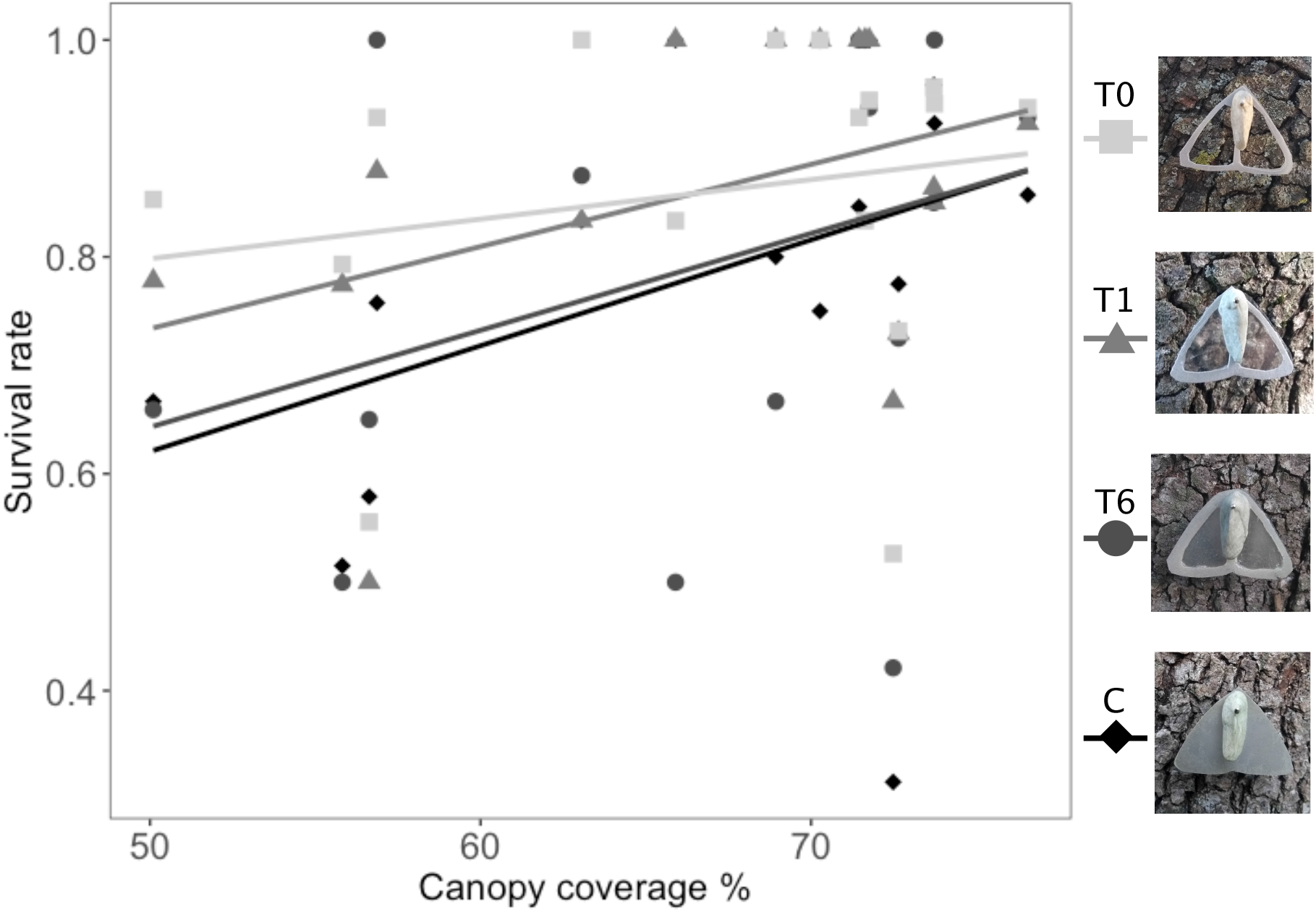
Survival rate of artificial prey per form according to the canopy coverage. Prey forms include: prey without any transparent area (C), and with transparent elements covered by 6 (T6), 1 (T1) or no (T0) transparent film layers. Open habitats exhibit low canopy coverage (towards the left of the plot) while closed habitats will have larger canopy cover (towards the right of the plot).

### 1.b. Experimental set up

Predation experiments were performed over four weeks between May and June 2018 at the zoological park of Montpellier (43.64°, 3.87°) and at La Rouvière forest (close to Montarnaud city 43.65°, 3.64°). Our experiment involved the local community of insectivorous birds and includes great tits (*Parus major*) and blue tits (*Cyanistes caeruleus*), which are reported visual predators in other similar studies (Rowland et al. 2008; Stevens et al. 2008a). Artificial prey items were pinned every 10 metres on oak trunks of at least 10 cm of diameter and with few or no moss cover. Prey order was randomized by blocks of eight items, two per treatment, before starting the experiment. Prey were mostly disposed facing north and as perpendicular to the floor as possible, to reduce direct sunlight reflection on them that could damage prey bodies with the heat and that could increase transparency shininess. Prey were checked every 24h. After 96h all prey were removed. When V- or U-marks were detected, or when the body was missing without signals of invertebrate attacks (i.e. no body scraps left on the wings or around the butterfly on the trunk) prey were considered as attacked and wings or pins where removed. Instead, if invertebrate attacks were detected, or when prey was missing, it was replaced. Non-attacked prey items were considered as censored data.

Canopy cover was used to assess habitat openness. Once the experiment was over, we took pictures each 20 meters in the trails that were used for the experiment (at forest edges for the open habitat, inside the forest for the closed habitat, examples Fig. S3), using a Canon EOS 550D camera and a fisheye lens. Canopy cover was further quantified using Fiji software (Schindelin et al. 2012), see Fig. S3 for the distribution of canopy cover in the experimental sites).

To test how different treatments survive along the experiment according to habitat openness, mixed effect and hierarchical (with block nested within location) Cox Proportional Hazards models (Cox 1972) were applied using the *coxme* package (Therneau 2020) in R (R Foundation for Statistical Computing 2014). Artificial type and habitat openness were included as explanatory variables. Models including or not their interaction were fitted and their fit was evaluated using the AIC criterion to select the best model.

### 2. Comparative analyses of museum specimens

### 2. a. Species selection and phylogeny

To explore the association of light transmittance with different conditions across multiple species we performed phylogenetic comparative analyses, since species are not independent. We used the phylogeny of Lepidoptera species exhibiting transparent areas in their wings by Gomez et al. (2021). In our analyses we included 107 species out of the 123 species included by Gomez et al, for which information about the openness of their habitat was available. These species spanned 31 of the 124 existing Lepidoptera families.

### 2. b. Ecological data on species habitat, activity rhythm and mimicry type

Ecological information was collected from the literature, Lepidoptera experts and our own knowledge for a total of 107 species for activity rhythm (57% of species are diurnal (day-active) species), habitat (56% of species living in open habitats), and mimicry status (14 species were bee/wasp mimics, 12 species were leaf mimics, while the remaining 81 did not fall in either of those categories (hereafter referred to as non-mimics although they include mimics involved in other butterfly mimetic communities, Table S1 for references and Fig S4). Habitat was considered open when it was described as “open habitat”, “meadow”, “small bushes”, “savannah”, “riversides”, or “rocky grasslands”. When the term “forested areas”, “primary forest”, “lowland forest”, “cloud forest” was mentioned the species was considered as closed habitat species (for references see Table S1). The selected 107 species excluded species that were reported to live in both open and closed habitats, or to fly at day and night (for references see Table S1). Visual predators are mostly active during the day and are supposed to be the main selective agents in the evolution of visual anti-predator defences (Ruxton et al. 2018). Characteristics that can increase conspicuousness for visual predators such as the presence of UV patterns (Lyytinen et al. 2004) and large body sizes (Guevara and Avilés 2013) are more commonly found in nocturnal than in diurnal insect species. Therefore, species that are active at night are unlikely to face similar selective pressures as species that are actively moving during the day although both could fly in similarly open habitats. For this reason, daytime rhythm and habitat were coded as a single variable with 4 states (diurnal open habitat, diurnal closed habitat, nocturnal open habitat and nocturnal closed habitat). To include mimicry syndrome, whenever species were described as bee and wasp mimics, they were assumed as so and they were grouped together in the same mimicry category, as both exhibit the striped pattern with black and often with yellow that advertise their unpalatability (Plowright and Owen 1980) and they exhibit similar body sizes. Butterflies and moths mimicking bees and wasps have forewings longer than their hindwings, and often strong delimitations of their wing borders and veins by dark coloration, as the hornet moth *Sesia apiformis* (*api - bee, formis* - shaped) or the broad-bordered bee hawk-moth *Hemaris fuciformis* (Fig 1b). Dead-leaf mimics were considered as so whenever they were described as death-leaf mimics on the literature and according to their general morphology. Dead-leaf mimics include brownish moths and butterflies with small or large transparent surfaces and that could touch or not the wing borders (Fig. 1a). *Rotschildia lebeau* has been proposed as a leaf mimic (Janzen 1984) but such mimicry is controversial (Hernández-Chavarría et al. 2004). *Rotschildia ericina* was therefore not considered as a leaf mimic for the analyses.

### 2. c. Wing structural and optical measurements

#### Macrostructure

Wing structural and optical measurements were taken from Gomez et al (2021), using the following methods. Museum specimens were photographed using a D800E Nikon camera equipped with a 60mm lens, placed on a stand with an annular light. Photos were then analysed using ImageJ (Schneider et al. 2012) to extract descriptors of wing macrostructure: wing length (mm), wing surface (mm^2^), and clearwing area (the surface of transparent area in mm^2^), for the forewing and hindwing separately. Proportion of clearwing area was considered by Gomez et al (2021) as the ratio clearwing area/wing surface, i.e. the proportion of the total wing area occupied by transparency. For optical measurements, Gomez et al (2021) followed the same procedure described above for measuring transmittance in artificial prey. For each species and wing, they took five measurements in the transparent zone, and analysed spectral shape using Avicol v6 (Gomez 2006) to extract the mean transmittance over [300-700] nm, which described the level of transparency.

#### Microstructure

Gomez et al (2021) described a large diversity in transparent wing surface characteristics. Using binocular imaging (Zeiss Stereo Discovery V20) and digital microscopic imaging (Keyence VHX-5000), these authors found that transparency could be achieved through the presence or absence of scales that had different morphologies, positions and colouration. Different combinations of structures and configurations are related to different light transmission levels. Therefore, we also explored the evolution of mimicry syndrome and of the different scale characteristics, including scale presence (nude membranes or with membranes covered by scales), type (lamellar scales, piliform scales or both), coloration (coloured or transparent) and insertion in the membrane (erect or flat).

### 2. e. Analyses of transmittance differences between habitats and mimics

To explore whether differences in the proportion of transmitted light are related to variation in habitat and mimicry we fitted both 1) a linear mixed model including 107 species for which we found data on habitat preference and 2) a Bayesian phylogenetic mixed model, including the phylogeny of the 107 species. Comparisons between the first and the second analysis allowed us to assess the influence, if any, of phylogenetic relationships on the observed trends. Light availability, mimicry and morphological traits were included as explanatory variables in our model. We defined *ActHab*, a variable combining activity rhythm and habitat openness with four levels (diurnal open, diurnal closed, nocturnal open, nocturnal closed) and three contrasts (diurnal vs. nocturnal, open vs. closed habitats, and diurnal open habitat vs. diurnal closed habitat). The variable mimicry has three levels (bee-wasp mimics, leaf mimics and non-mimics) and we tested two contrasts (bee/wasp mimics vs. all other species, and leaf mimics vs. non-mimics). Other explanatory variables included: wing length (as a proxy of butterfly size), proportion of clearwing surface and fore-/hindwing, reported by Gomez et al (2020) as correlated to light transmittance. Wing transmittance, wing length and clearwing proportion were obtained from Gomez *et al* (2021). As five measurements per wing (fore-and hindwing), thus ten measurements per species were included in our dataset, wing measurement replicates nested in species was considered as a random effect. We fitted different models including different combinations of two-factor interactions and we compared them using the AIC criterion to select the best model. As mimicry levels show differences in the variance of their transparency degree (see results for Fligner test and Fig. 4), we estimated different variances for the different factor levels by specifying a variance structure into the model, using the function *varIdent*. To better understand the effect of each of the included variables, we reported and compared the full model without interactions and the best model including interactions (Tables S1 and S3). The Bayesian phylogenetic mixed models with Markov chain Monte Carlo analyses were performed using the ‘mulTree’ R package (Guillerme and Healy 2014). Using the best formulated linear mixed model, uninformative priors were used with an Inverse-Gamma distribution with shape = 0.001 and scale = 0.001 for both random effect and residual variances (Hadfield 2010). Models were run using two chains of 600,000 iterations, a thinning interval of 300 and a burn-in of 10,000. Fixed effects were considered statistically significant when the probabilities in the 95% credible intervals did not include zero. As Bayesian phylogenetic mixed models consider no interactions between factor levels, several submodels were fitted to test the interactions from the best model obtained according to AIC criterion.

We also explored the coevolution between other factors that are correlated to conspicuousness, transparency degree and mimicry. At least in the tropics, insects that are active during the day are smaller, on average, than insects active at night a segregation probably related to the higher diurnal visual predation risk and the higher detectability associated to larger bodies (Guevara and Avilés 2013). Moreover, Gomez et al (2021) showed that transmittance was higher at smaller wing sizes and larger proportions of clearwing surface. Therefore, we performed several phylogenetic generalised least square (PGLS) analyses (1 observation per species) to test whether small wing size and large clearwing surface proportion associate with diurnality, open habitats and/or bee/wasp mimicry. Additionally, using also PGLS we tested whether wing ratio (forewing length/hindwing length), which is usually high in bees and wasps, is higher in bee/wasp mimics than dead-leaf mimics or in non-mimics. Finally, using BayesTraits (Pagel and Meade 2013) we tested whether bee/wasp mimics are more often diurnal and/or live in more open areas than non- or dead-leaf mimics. For this last test, we calculated the likelihood of the dependent (correlated) and independent (non-correlated) models of evolution mimicry syndrome and *ActHab* levels using a Markov-Chain Monte-Carlo approach. Models were compared using the likelihood ratio test with 4 degrees of freedom (Pagel 1994). As this approach can only be applied to binary characters, and mimicry has three factors, models were run with different pairwise combinations of variables to explore: a. the evolution of dead-leaf mimicry (including non-mimics and dead-leaf mimics); b. evolution of bee/wasp mimics (including non-mimics and bee/wasp mimics) and c. differences in the evolution of dead-leaf and bee/wasp-mimics (including bee/wasp and dead-leaf mimics). Similarly, for the composed variable *ActHab* we compared (i) diurnal and nocturnal species flying in open habitats, and (ii) nocturnal species flying in open and in closed habitats.

We also tested whether scale characteristics associated to differences in light transmittance (as described by Gomez et al (2021)) have coevolved with mimicry syndrome and their specific optical properties, using again the test of correlated evolution implemented in BayesTraits (Pagel and Meade 2013). Scale characteristics included in this analyses were: absence/presence of scales, scale insertion (flat or erect), scale coloration (coloured or transparent) and scale type (piliform scales and lamellar scales). Independent analyses were performed for anterior and posterior wings, as differences between wings have been reported by Gomez et al (2021).

Finally, we predicted higher variance in light transmittance for those conditions where different levels of transparency might be equally efficient, as would be the case for butterflies and moths that are active during the night, that live in closed habitats and/or that are dead-leaf mimics. We applied Fligner-Killeen tests of homogeneity of variance in R to test for difference in variance between (i) diurnal and nocturnal species living in open and in close habitats, and (ii) bee/wasp mimics, leaf mimics and no mimic species. As Gomez et al (2021) reported that transmittance is closely related to wing size, we also tested for differences in the variation of wing length between habitat, activity rhythm and mimicry using the test mentioned above.

## Results

### Fieldwork experiment

A total of 243 out of the 1149 artificial butterflies were attacked (21.14 %). The best model included artificial type and habitat openness without interaction (Delta AIC 2.79 between a model including and excluding the interaction). Artificial moth survival was lower in more open habitats (*z* = −2.67, p = 0.007, Figure 2). Fully opaque moths “C” and the least transparent moths (T6) show similar survival (Figure 2) and together were more attacked than both T1 and T0 (*z* = 3, p = 0.003). However, no differences are detected anymore when comparing survival of T6 and either T0 (*z* = 0.44, p = 0.66) or T1 (*z* = 1.25, p = 0.21).

### Comparative analyses

### Relationships between light transmittance, habitat and activity rhythm

Activity rhythm and habitat openness partly explain the variation in light transmission. Diurnal and nocturnal species transmit light in a similar way (Estimate and confidence interval for Bayesian models without interaction: 0.46 [-10.07, 10.82]; with interaction −5.2 [−18.52, 7.72], Figures 3a & 5, Table S1 & S2a). However, diurnal (D) species transmit slightly more light than nocturnal (N) species at smaller clearwing surfaces and at larger wing lengths (D>N: %clearwing surface: −0.18 [−0.28, −0.18]; D>N: wing length 0.46[0.13,0.77], Figures 5, S4a & S4b, Table S1 & S2a). Moreover, forewings (F) transmit more light in diurnal than in nocturnal species (D>N:F>H : 8.41[4.76, 12.12]). Wing length and proportion of clearwing surface is similar in diurnal and nocturnal species (Table S2a). Variation in forewing light transmission is lower for diurnal than for nocturnal species (Fligner-Killeen test for forewings: 17.44, df=1, p< 0.001, Figure 3a). Similarly, wing size variation is lower in diurnal than in nocturnal species and for both fore- and hindwings (Fligner-Killeen test for forewings: 8.87, df=1, p= 0.003; for hindwings: 28.86, df=1, p < 0.001, Figure 3b).

**Figure 3:**
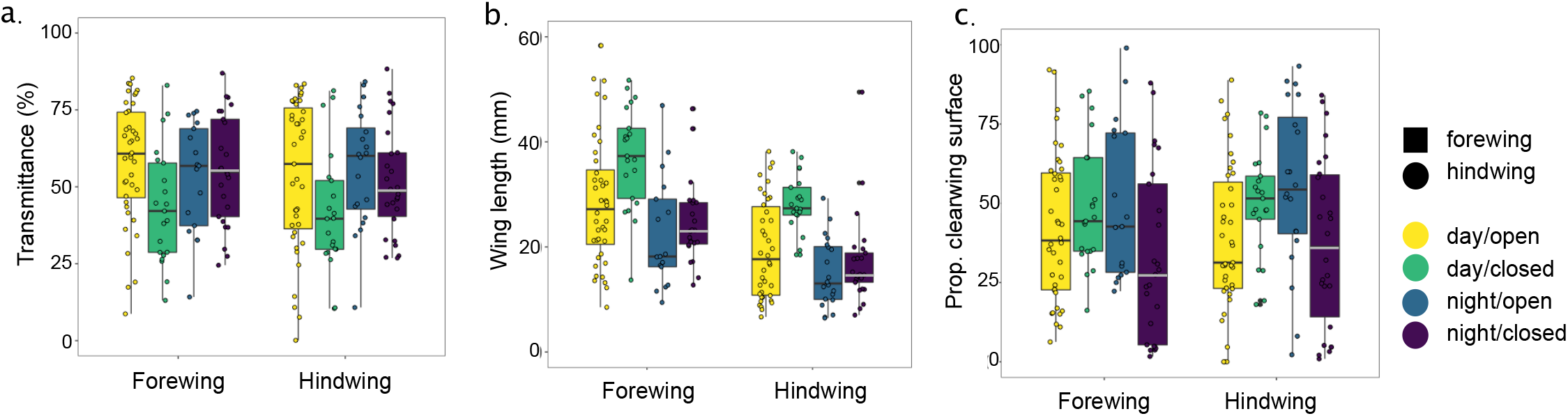
Variation between diurnal/open (yellow), diurnal/closed (green), nocturnal/open (blue) and nocturnal/closed (purple) in a) mean light transmittance, b) wing length and c) proportion of clearwing surface. Values larger than 60 mm were not plotted for clarity reasons but were included in the analyses.

**Figure 4:**
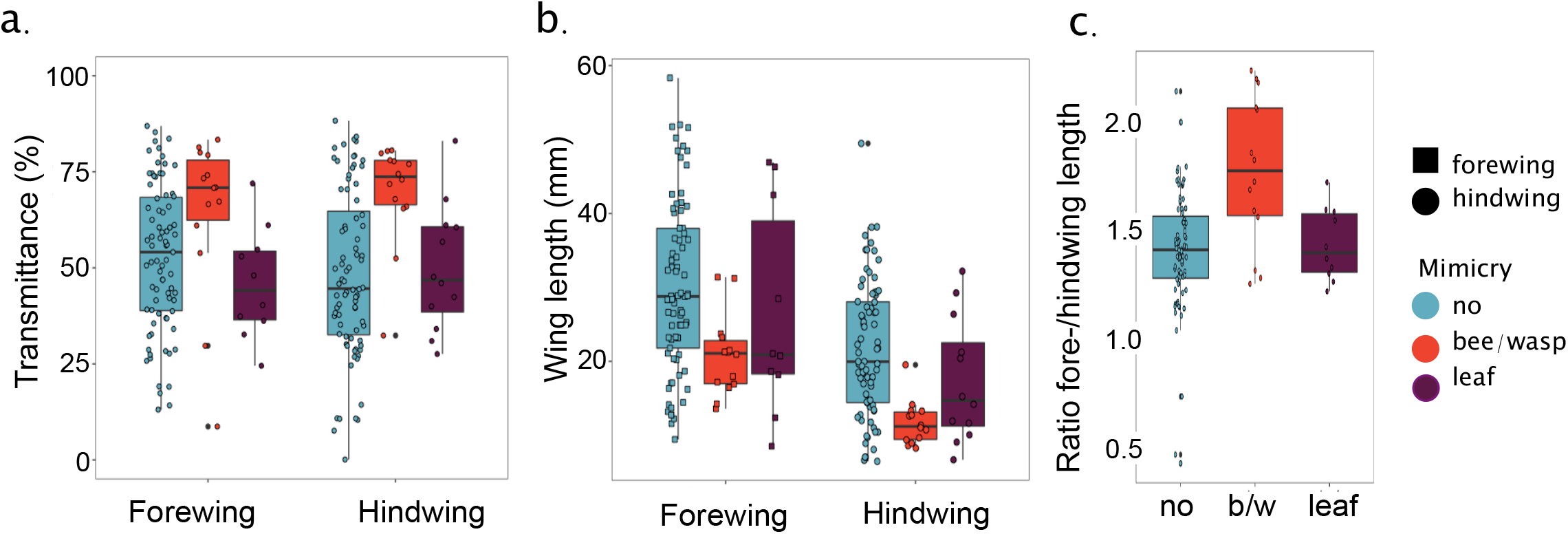
Variation between bee/wasp mimics (red), leaf mimics (purple) and no mimics (blue) in a) mean light transmittance, b) wing length and c) ratio between length of forewing and hindwing per mimicry group:

**Figure 5.**
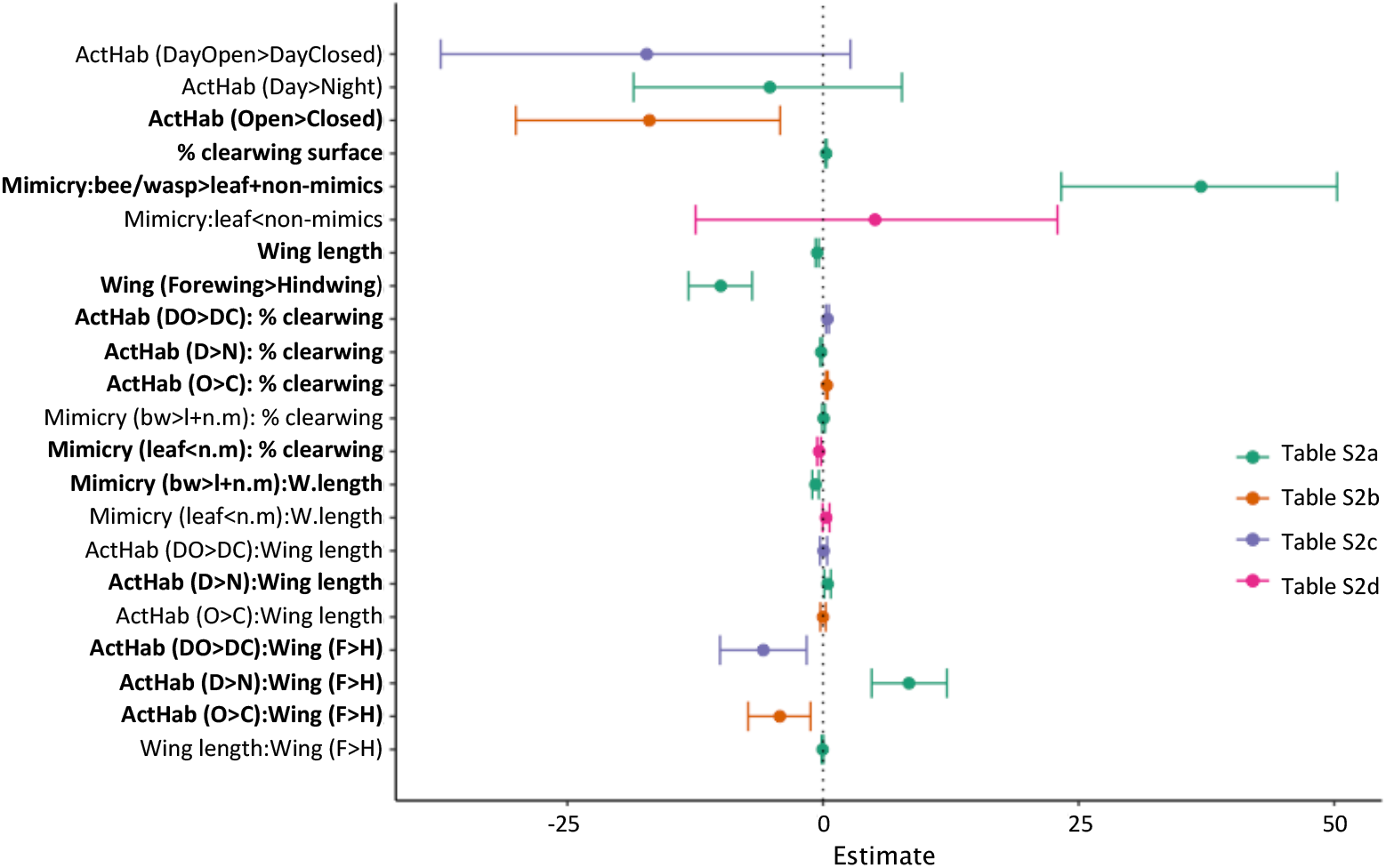
Bayesian regression estimates and 95% confidence intervals for each variable and their interactions in the best fitted model. Such model included transmittance as dependent variable and the following explanatory variables: the combination between daytime activity and habitat type (ActHab: nocturnal-open/nocturnal-closed/diurnal-open/diurnal-closed), wing length in mm, proportion of clearwing area (% clearwing), mimicry syndrome (bee/wasp-, leaf- or non-mimics), wing (fore-/hindwing) and the interactions: mimicry syndrome and proportion of clearwing area, mimicry syndrome and wing size, proportion of clearwing area and habitat type, wing size and habitat type, wing size and proportion of clearing area, wing and wing length and proportion of clearwing area, wing length and habitat as explanatory variables. Wing measurements nested in species was considered as random effect. Names of variables and interactions whose confidence interval excludes zero are in bold. Model results are found in Tables S2.

Regarding habitat openness, we found that light transmittance is slightly higher for species living in closed than in open habitats (with interactions: −16.98 [−30.04, −4.19], Table S2b). However, two characteristics associated to higher light transmission are more common in open habitats: the proportion of clearwing surface is larger for species flying in open than in closed habitats (PGLS forewing: t = 4.02, p <0.001; hindwing: t= 2.01, p = 0.04, Table S3) and butterflies and moths flying in open habitats have smaller forewings than species flying in closed habitats (PGLS forewing: t = −2.12, p = 0.04, Table S3). When the proportion of clearwing surfaces is high, species flying in open habitats (O) transmit slightly more light than species living in closed habitats (C) (O>C:%clearwing surface: 0.44 [0.3,0.59], Figure S4a). Variation in light transmission and wing length are similar between open and closed habitats (Fligner-Killeen test for light transmission in forewings: 3.23, df=1, p= 0.07; in hindwings: 1.66, df=1, p = 0.20. Fligner-Killeen test for length in forewings: 0.24, df=1, p= 0.62; in hindwings: 0.19, df=1, p = 0.66, Figures 3b & 3c)

When analysing only diurnal species, we found that light transmittance is similar between species flying in open and closed habitats (without interactions: 1.64 [-8.88,12.88], with interactions: −17.24 [−37.39,2.68], Table S2c). However, at larger proportions of clearwing surfaces, diurnal species flying in open habitats transmit slightly more light than diurnal species flying in closed habitats (DO>DC:%clearwing surface: 0.44 [0.3,0.58], Figure 3c & 5, Table S1 & S2c). Nevertheless, diurnal species flying in closed habitats exhibit larger proportions of clearwing surface than do species flying in open habitats (PGLS t=−4.34, p<0.001, Table S3). Transmittance in the hindwing of diurnal species flying in closed habitats varies less than for species flying in open habitats (Fligner-Killeen test for forewing: 0.06, df=1, p= 0.80, hindwing: 6.45, df=1, p=0.01, Figure 3a). In diurnal species, open habitat species exhibit larger variation in wing length than closed habitat species (Fligner-Killeen test for forewing: 5.9, df=1, p=0.015, hindwing: 43.91, df=1, p<0.001, Figure 3b).

### Relationships between light transmittance, transparency structural basis and mimicry

Mimicry also explains part of the variation found in light transmittance. Bee/wasp mimics transmit more light than dead-leaf mimics and other clearwing species (without interactions: 26.38 [15.16, 37.12]; with interactions: 36.97 [23.3, 50.28], Figures 4a & 5, Tables S1 & S2a). Light transmittance in bee/wasp mimics is slightly lower at larger wing lengths (bee/wasp mimics>others species: wing length: −0.73 [−1.04, −0.41]; Figure 5, Tables S1 & S2a). Additionally, non-mimics transmit slightly less light than dead-leaf mimics at larger proportions of transparent wing surface (leaf mimics<non-mimics: %clearwing surface: −0.37 [−0.57, −0.19]; Figures 5 & S5b, Tables S1 & S2d).

We found that transmittance varies more in dead-leaf mimics than in bee/wasp mimics (Fligner-Killeen test for forewing: 31.38, df= 1, p<0.001, for hindwing: 50.43, df = 1, p<0.001), or in other clearwing species (Fligner-Killeen test for forewing: 37.95, df=1, p<0.001; for hindwing: 57.51, df = 1, p<0.001), but we found no difference in transmittance variation between leaf mimics and non-mimic species (Fligner-Killeen test for forewing: 2.02, df=1, p= 0.155; for hindwing: 0.00005, df = 1, p=0.995). Likewise, wing length varies less in bee/wasp mimics than in leaf mimics (Fligner-Killeen test for forewing: 16.96, df=1, p<0.001; for hindwing: 37.98, df = 1, p<0.001, Figure 4b) or in other clearwing species (Fligner-Killeen test for forewing: 37.87, df=1, p<0.001; for hindwing: 57.39, df = 1, p<0.001). Leaf and non-mimics show similar variations in wing length (Fligner-Killeen test for forewing: 0.68, df=1, p=0.40; for hindwing: 2.52, df = 1, p=0.11).

Regarding the relationship between mimicry and wing characteristics, non-mimics have smaller hindwings than dead-leaf mimics (PGLS forewing: t=1.82, p = 0.07; hindwing: t=1.99, p = 0.05, Table S3), and the length ratio forewing/hindwing is larger for bee/wasp mimics than for any other clearwing species group (PGLS t=4.14, p <0.001, Fig. 4c). Additionally, the proportion of clearwing surface in forewings and hindwings is larger for non-mimics and for dead-leaf mimics (PGLS forewing: t=2.74, p=0.007; hindwing: t=2.29, p=0.02, Figure S5a). Mimicry syndrome and activity rhythm have likely coevolved: most bee/wasp mimics are diurnal while most leaf mimics are nocturnal (BayesTraits activity rhythm and bee/wasp or leaf mimics: LRT=11.43, df = 4, p=0.02, Table 2, Figure S3 & S6).

Some scale characteristics are correlated to mimicry evolution. Scales are more often absent and less often coloured and flat when present in hindwings of bee/wasp mimics in comparison to dead-leaf mimics (BT LRT presence = 10.45, df=4, p=0.033; BT LRT insertion= 11.06, df=4, p=0.03) and non-mimics wings (BT LRT presence = 29.99, df=4, p=0.001; BT LRT colour= 21.34, df=4, p=0.001; BT LRT insertion = 11.17, df=4, p=0.025, Table S4 & Fig S7). Similarly, dead-leaf mimics have less often coloured scales in comparison to non-mimics both in forewing (BT LRT=11.62, df=4, p=0.02) and hindwing (BT LRT=12.87, df=4, p=0.012, Table S4 & Fig S7). Moreover, leaf mimics only exhibit lamellar scales in contrast to non-mimics (BT LRT=9.33, df=4, p=0.053, Table S4 & Fig S7).

## Discussion

Fieldwork experiments using artificial prey that differed in their proportion of transmitted light suggest that variation in transparency is similarly detected in bright and dim light conditions, contrasting with what has been suggested for aquatic environments (Johnsen and Widder 1998). Opaque prey were more attacked than highly transparent prey in both open and closed habitats, suggesting that transparency enhances crypsis in terrestrial environments, independently of light conditions. Moreover, T6 survival is marginally the lowest in comparison to the other transparent types (T1 and T0), suggesting that terrestrial predators probably detect differences in transparency degree as well as aquatic predators do. The lack of larger differences between artificial prey types might be related to the continuum created by our moths that could have elicited only slightly different reactions towards each of them as in other similar studies (Arias et al 2021). An experiment where higher number of responses towards each type is be obtained, can contribute to the understanding of this trend. Moreover, whether factors such as movement can enhance differences between transparency degrees remains to be tested. Additionally, fewer artificial moths were attacked in closed habitats, in accordance to the higher predation rate often reported for open habitats such as forest edges (Barbaro et al. 2014; Seymoure et al. 2018). Such higher predation rate can be related to a higher detectability of prey in such exposed areas (Stankowich and Campbell 2016) and in the case of forest edges, to their use as travel lanes by predators, increasing their presence in those places (Andrén 1995; Saab 1999).

Our comparative analyses show some evidence supporting that variation in transparency degree is related to light availability. Diurnal species transmit more light on their forewings and exhibit a higher transparency degree at lower percentages of clearwing surface and at larger wing length sizes than nocturnal species. Moreover, diurnal species vary less in the light they transmit than nocturnal species, suggesting a strong selection for high levels of transmittance. Although both diurnal and nocturnal species are exposed to visual predators during the day, species that are active during the day, independently of the habitat where they fly, are at higher predation risk as they can be more easily detected by their movement (Lyytinen et al. 2004; Galloway et al. 2020). Therefore, traits reducing this detectability, such as high transparency levels will be strongly selected.

However and in accordance to our experimental results, no differences in transmittance were detected between diurnal species flying in open and closed habitats. Additionally, butterflies and moths living in closed habitats (including both diurnal and nocturnal) transmit similar to slightly more light than species living in open habitats. Gomez et al (2021) reported that poor light transmittance is associated with small proportions of clearwing surface and large wing sizes. Diurnal species living in closed habitats show higher proportion of clearwing surface than diurnal species living in open habitats (Table 1, Fig. 3), probably contributing to the similar/slightly higher values of light transmittance in closed habitats in comparison to the open habitats. Selective pressures related to abiotic conditions might also help understanding this pattern. Species living in open habitats undergo higher risks of tissue damage by higher exposure to UV light, more wind or higher exposure to direct rainfall, conditions that have been reported as important at shaping moth and butterfly segregation in different habitats (Brown Jr and Hutchings 1997). In case transparency entails costs related to one or several of these factors, selection may promote lower proportions of clearwing surfaces in open habitats, decreasing the vulnerability of clearwing Lepidoptera species to these abiotic effects, and light transmittance by default.

**Table 1.**
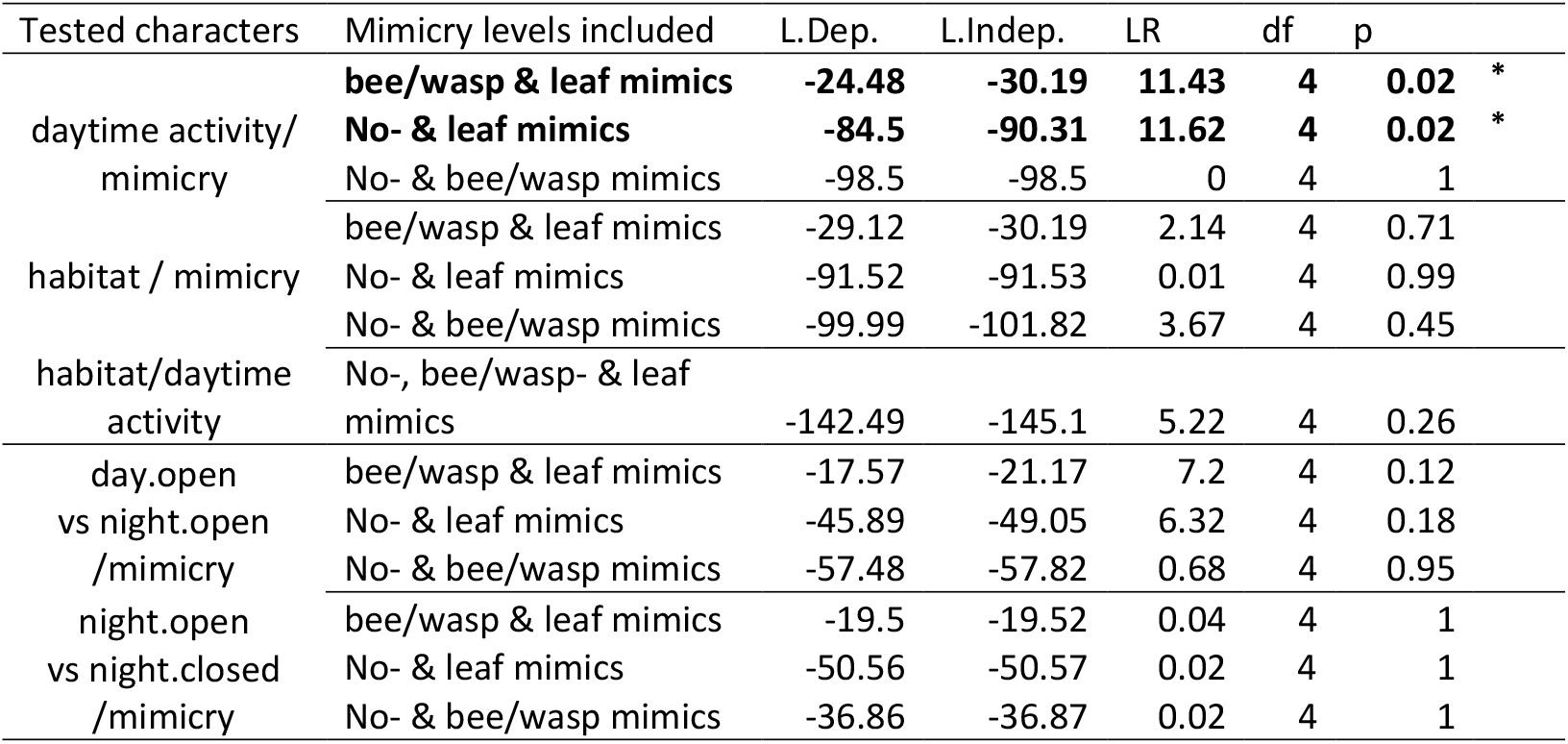
Results of coevolution tests between habitat, mimicry and daytime activity Log-likelihood of dependent (L. Dep. Coevolution, 4 parameters estimated) and independent (L. Indep. 8 parameters estimated) evolution models between pairs of binary characters obtained from BayesTraits, log-likelihood ratio (LR), associated degree of freedom (df: difference in number of parameters) and corresponding p-value of the Likelihood Ratio Test. Because BayesTraits only take binary characters, when factors have more than 2 levels (mimicry and the composed character ActHab), we implemented pairwise comparisons of level. Coevolution between mimicry and diurnal species flying either in open or in closed habitats was not tested as no bee/wasp mimic is simultaneously diurnal and closed habitat dweller (Fig S4). A significant p-value means that the dependent model is significantly better than the independent model, indicating a correlated evolution between the traits. ** stands for statistical significance below 0.01 and * for significance values below 0.05.

Transparency can decrease prey detectability but it is also involved in mimicry of bees or wasps, and differences in light transmittance seem to be closely associated to mimicry evolution. We found that bee/wasp mimics are diurnal (similar to most bees, Somanathan et al. 2009), and have smaller wing sizes and smaller hindwings in comparison to their forewings (as has been reported for wasps and bees, Wootton 1992; Lindauer 2019). Additionally, they have higher transmittance levels, frequently associated to the absence of scales in their wings, and are more often restricted to open habitats suggesting that high transmittance in open habitats can be a visual optimum in terrestrial environments under certain circumstances. As this is not the case for all clearwing species, this highlights the strong effect of biotic and abiotic interactions in shaping adaptive peaks, thus the segregation of species under different conditions and the evolution of different features. However, the correlation between open habitats, diurnal activity, small sizes and higher transmittance levels, suggests that transparency in bees, wasps and their mimics probably reduces their detectability, a question that remains open.

On the other hand, we found that leaf mimics are more often found in closed habitats, as expected given to the higher presence of dead leafs in closed habitats. Additionally, they are larger and in contrast to the trends for all other clearwing species, transmit more light at lower proportions of clearwing surfaces. Some fully opaque fallen leaf mimic species, such as the comma butterfly (*Polygonia c-album*) exhibit a distinctive pale mark that might decrease their predation by being perceived as ‘distractive marks’ preventing predators from recognising prey, thus enhancing background matching (Dimitrova et al. 2009; Olofsson et al. 2013), although it is controversial (Troscianko et al. 2018). Either way, reduced and highly transparent zones as those found in *Orthogonioptilum violascens* females might produce a similar appearance to those markings; their high transmittance might be highly reflective and be perceived as a white mark, thus conferring the potential benefits of the “comma mark” when seen for a given angle, as well as the full crypsis, when no reflections can be perceived. Whether highly transparent windows work as other highly visible elements displayed in cryptic species such as eyespots that deter predators (Stevens et al. 2008b, 2009) and that are also important in mate choice (Robertson and Monteiro 2005) remains to be tested.

Other dead-leaf mimics include large transparent surfaces that can expand to the wings edges, as for *Bertholdia* species, or not, as for *Pseudasellodes fenestraria*. In both cases, transparent surfaces likely disrupt prey shape (as in the first case) and/or prey surface (as in the second example (Costello et al. 2020)), thereby reducing their detectability. Disruptive coloration, especially when touching wing margins, is common in fully opaque species with cryptic patterns/colours and has been broadly studied as it is common in fully opaque cryptic species (Cuthill et al. 2005; Schaefer and Stobbe 2006; Stevens and Cuthill 2006; Fraser et al. 2007). Whether transparent surfaces touching wing margins, work as disruptive marks regardless of their transmittance, thus further decreasing prey detectability remains to be tested.

## Conclusion

Both light availability (associated to activity rhythm and habitat openness) and mimicry can concur to fostering variations in light transmittance on land. Our results opens up new questions that need to be investigated in the future, in particular the effect of both biotic (such as mimicry interactions with different models) and abiotic (such as the possible cost associated to the physical conditions of each habitat), to better understand the evolution and the constraints imposed on transparency on land.

## Supporting information

ESM

## References

Andrén, H. 1995. Effects of landscape composition on predation rates at habitat edges. Pages 225–255 inMosaic landscapes and ecological processes. Springer.

Arias, M., M. Elias, C. Andraud, S. Berthier, and D. Gomez. 2020. Transparency improves concealment in cryptically coloured moths. Journal of Evolutionary Biology 33:247–252.

Arias, M., J. Mappes, C. Desbois, S. Gordon, M. McClure, M. Elias, O. Nokelainen, et al. 2019. Transparency reduces predator detection in mimetic clearwing butterflies. Functional Ecology 33:1110–1119.

Barbaro, L., B. Giffard, Y. Charbonnier, I. van Halder, and E. G. Brockerhoff. 2014. Bird functional diversity enhances insectivory at forest edges: a transcontinental experiment. Diversity and Distributions 20:149–159.

Bates, H. W. 1862. XXXII. Contributions to an Insect Fauna of the Amazon Valley. Lepidoptera: Heliconidæ. Transactions of the Linnean Society of London 23:495–566.

Brown Jr, K. S., and R. W. Hutchings. 1997. Disturbance, fragmentation, and the dynamics of diversity in Amazonian forest butterflies. Tropical forest remnants: ecology, management, and conservation of fragmented communities. University of Chicago Press, Chicago 632.

Costello, L. M., N. E. Scott-Samuel, K. Kjernsmo, and I. C. Cuthill. 2020. False holes as camouflage. Proceedings of the Royal Society B 287:20200126.

Cox, D. R. 1972. Models and life-tables regression. JR Stat. Soc. Ser. B 34:187–220.

Cuthill, I. C., M. Stevens, J. Sheppard, T. Maddocks, C. A. Párraga, and T. S. Troscianko. 2005. Disruptive coloration and background pattern matching. Nature 434:72.

Dimitrova, M., N. Stobbe, H. M. Schaefer, and S. Merilaita. 2009. Concealed by conspicuousness: distractive prey markings and backgrounds. Proceedings of the Royal Society B: Biological Sciences 276:1905–1910.

Fraser, S., A. Callahan, D. Klassen, and T. N. Sherratt. 2007. Empirical tests of the role of disruptive coloration in reducing detectability. Proceedings of the Royal Society of London B: Biological Sciences 274:1325–1331.

Galloway, J. A., S. D. Green, M. Stevens, and L. A. Kelley. 2020. Finding a signal hidden among noise: how can predators overcome camouflage strategies? Philosophical Transactions of the Royal Society B 375:20190478.

Gomez, D. 2006. AVICOL, a program to analyse spectrometric data. Free executable available at http://sites.google.com/site/avicolprogram/ or from the author at. Last update october 2011.

Gomez, D., C. Pinna, J. Pairraire, M. Arias, J. Barbut, A. Pomerantz, C. Noûs, et al. 2021. Transparency in butterflies and moths: structural diversity, optical properties and ecological relevance. Ecological Monogrpahs 2020.05.14.093450.

Grundel, R., R. P. Jean, K. J. Frohnapple, G. A. Glowacki, P. E. Scott, and N. B. Pavlovic. 2010. Floral and nesting resources, habitat structure, and fire influence bee distribution across an open-forest gradient. Ecological applications 20:1678–1692.

Guevara, J., and L. Avilés. 2013. Community-wide body size differences between nocturnal and diurnal insects. Ecology 94:537–543.

Guillerme, T., and K. Healy. 2014. mulTree: a package for running MCMCglmm analysis on multiple trees. Zenodo. (doi: 10.5281/zenodo.12902).

Hadfield, J. D. 2010. MCMC Methods for Multi-Response Generalized Linear Mixed Models: The MCMCglmm R Package. Journal of Statistical Software 33.

Hernández-Chavarría, F., A. Hernández, and A. Sittenfeld. 2004. The” windows”, scales, and bristles of the tropical moth Rothschildia lebeau (Lepidoptera: Saturniidae). Revista de biología tropical 52:919–926.

Hodos, W., R. W. Leibowitz, and J. C. Bonbright Jr. 1976. Near-field visual acuity of pigeons: Effects of head location and stimulus luminance. Journal of the Experimental Analysis of Behavior 25:129–141.

Janzen, D. H. 1984. Weather-related color polymorphism of Rothschildia lebeau (Saturniidae). Bulletin of the ESA 30:16–21.

Johnsen, S. 2014. Hide and seek in the open sea: pelagic camouflage and visual countermeasures. Annual review of marine science 6:369–392.

Johnsen, S., and E. A. Widder. 1998. Transparency and visibility of gelatinous zooplankton from the northwestern Atlantic and Gulf of Mexico. The Biological Bulletin 195:337–348.

Kassarov, L. 2003. Are birds the primary selective force leading to evolution of mimicry and aposematism in butterflies? An opposing point of view. Behaviour 140:433–451.

Lind, O., S. Karlsson, and A. Kelber. 2013. Brightness discrimination in budgerigars (Melopsittacus undulatus). PLoS One 8:e54650.

Lindauer, M. 2019. Hymenopteran. Encyclopædia Britannica. Encyclopædia Britannica, inc.

Lyytinen, A., L. Lindström, and J. Mappes. 2004. Ultraviolet reflection and predation risk in diurnal and nocturnal Lepidoptera. Behavioral Ecology 15:982–987.

Mcclure, M., C. Clerc, C. Desbois, A. Meichanetzoglou, M. Cau, L. Bastin-Héline, J. Bacigalupo, et al. 2019. Why has transparency evolved in aposematic butterflies? Insights from the largest radiation of aposematic butterflies, the Ithomiini. Proceedings of the Royal Society B 286:20182769.

Olofsson, M., M. Dimitrova, and C. Wiklund. 2013. The white ‘comma’as a distractive mark on the wings of comma butterflies. Animal behaviour 86:1325–1331.

Pagel, M. 1994. Detecting correlated evolution on phylogenies: a general method for the comparative analysis of discrete characters. Proceedings of the Royal Society of London. Series B: Biological Sciences 255:37–45.

Pagel, M., and A. Meade. 2013. Bayes Traits V2. Computer program and documentation. Available at: http://www.evolution.rdg.ac.uk/BayesTraits.html (accessed 12 July 2013).

Pinna, C., M. Vilbert, S. Borenztajn, W. D. de Marcillac, F. Piron-Prunier, A. Pomerantz, N. Patel, et al. 2021. Mimicry drives convergence in structural and light transmission features of transparent wings in Lepidoptera. bioRxiv 2020.06.30.180612.

Plowright, R., and R. E. Owen. 1980. The evolutionary significance of bumble bee color patterns: a mimetic interpretation. Evolution 622–637.

Pomerantz, A., R. Siddique, E. Cash, Y. Kishi, C. Pinna, K. Hammar, D. Gomez, et al. 2021. Developmental, cellular, and biochemical basis of transparency in the glasswing butterfly Greta oto.

R Foundation for Statistical Computing, R. C. 2014. R: A language and environment for statistical computing. Vienna, Austria.

Robertson, K. A., and A. Monteiro. 2005. Female Bicyclus anynana butterflies choose males on the basis of their dorsal UV-reflective eyespot pupils. Proceedings of the Royal Society of London Series B-Biological Sciences 272:1541–1546.

Rowland, H. M., I. C. Cuthill, I. F. Harvey, M. P. Speed, and G. D. Ruxton. 2008. Can’t tell the caterpillars from the trees: countershading enhances survival in a woodland. Proceedings of the Royal Society B: Biological Sciences 275:2539–2545.

Ruxton, G. D. Allen, William L., T. N. Sherratt, and M. P. Speed. 2018. Avoiding attack: the evolutionary ecology of crypsis, warning signals and mimicry (Second.). Oxford University Press.

Saab, V. 1999. Importance of spatial scale to habitat use by breeding birds in riparian forests: a hierarchical analysis. Ecological applications 9:135–151.

Schaefer, H. M., and N. Stobbe. 2006. Disruptive coloration provides camouflage independent of background matching. Proceedings of the Royal Society of London B: Biological Sciences 273:2427–2432.

Schindelin, J., I. Arganda-Carreras, E. Frise, V. Kaynig, M. Longair, T. Pietzsch, S. Preibisch, et al. 2012. Fiji: an open-source platform for biological-image analysis. Nature methods 9:676.

Schneider, C. A., W. S. Rasband, and K. W. Eliceiri. 2012. NIH Image to ImageJ: 25 years of image analysis. Nature Methods 9:671–675.

Seymoure, B. M., A. Raymundo, K. J. McGraw, W. Owen McMillan, and R. L. Rutowski. 2018. Environment-dependent attack rates of cryptic and aposematic butterflies. Current zoology 64:663–669.

Skelhorn, J., H. M. Rowland, M. P. Speed, and G. D. Ruxton. 2010. Masquerade: camouflage without crypsis. Science 327:51–51.

Somanathan, H., A. Kelber, R. M. Borges, R. Wallén, and E. J. Warrant. 2009. Visual ecology of Indian carpenter bees II: adaptations of eyes and ocelli to nocturnal and diurnal lifestyles. Journal of Comparative Physiology A 195:571–583.

Stankowich, T., and L. A. Campbell. 2016. Living in the danger zone: exposure to predators and the evolution of spines and body armor in mammals. Evolution 70:1501–1511.

Stevens, M., A. Cantor, J. Graham, and W. I.S. 2009. The function of animal “eyespots”: conspicuousness but not eye mimicry is key. Current Zoology 55:319–326.

Stevens, M., and I. C. Cuthill. 2006. Disruptive coloration, crypsis and edge detection in early visual processing. Proceedings of the Royal Society B: Biological Sciences 273:2141.

Stevens, M., C. J. Hardman, and C. L. Stubbins. 2008a. Conspicuousness, not eye mimicry, makes “eyespots” effective antipredator signals. Behavioral Ecology 19:525–531.

Stevens, M., C. L. Stubbins, and C. J. Hardman. 2008b. The anti-predator function of “eyespots” on camouflaged and conspicuous prey. Behavioral Ecology and Sociobiology 62:1787–1793.

Therneau, T. M. 2020. Mixed Effects Cox Models [R package coxme version 2.2-16].

Troscianko, J., J. Skelhorn, and M. Stevens. 2018. Camouflage strategies interfere differently with observer search images. Proceedings of the Royal Society B: Biological Sciences 285:20181386.

Wootton, R. J. 1992. Functional morphology of insect wings. Annual review of entomology 37:113–140.

Yamaura, Y., J. A. Royle, N. Shimada, S. Asanuma, T. Sato, H. Taki, and S. Makino. 2012. Biodiversity of man-made open habitats in an underused country: a class of multispecies abundance models for count data. Biodiversity and Conservation 21:1365–1380.

