## Supplementary material for "Ambient light and mimicry as drivers of wing transparency in Lepidoptera": ESM

**Electronic supplementary material- Ambient light and mimicry as drivers of wing transparency in Lepidoptera**

Table S1. Information for the 107 clearwing species included on this study.

| Species | Family | Subfamily | Tribe | Activity | Habitat | Mimicry | Information sources |
| --- | --- | --- | --- | --- | --- | --- | --- |
| *Colla netrix* | Bombycidae | Epiinae | na | N | C | no | Expert R. Rougerie |
| *Castnia heliconioides* | Castniidae | Castniinae | na | D | O | no | (Worthy et al. 2019) |
| *Eulophonotus myrmeleon* | Cossidae | Zeuzerinae | na | D | O | b/w | (N’guessan et al. 2010) |
| *Zeuzera pyrina* | Cossidae | Zeuzerinae | na | N | O | no | Expert J. Barbut |
| *Diaphania unionalis* | Crambidae | Spilomelinae | na | N | O | no | (Negm 1968) |
| *Glyphodes pyloalis* | Crambidae | Spilomelinae | na | N | O | leaf | (Park et al. 2016) |
| *Minacraga hyalina* | Dalceridae | na | na | N | C | no | (Miller 1994) |
| *Anadasmus sp* | Depressariidae | Stenomatinae | na | N | O | no | Other species from the subfamily - (Urra 2014) |
| *Ethmia aurifluella* | Depressariidae | Ethmiinae | na | N | O | no | (Nupponen 2015) |
| *Macrauzata limpidata* | Drepanidae | Drepaninae | na | N | C | no | (Holloway 1999) |
| *Aethria andromacha* | Erebidae | Arctiinae | Ctenuchini | N | C | b/w | Expert J. Barbut |
| *Amastus parergana* | Erebidae | Arctiinae | Phaegopterini | N | O | no | Expert J. Barbut |
| *Amata mogadorensis* | Erebidae | Arctiinae | Syntomini | D | O | no | (Sbordoni et al. 1979) |
| *Bertholdia sp* | Erebidae | Arctiinae | Arctiini | N | C | leaf | (Beccacece 2017) |
| *Carriola ecnomoda* | Erebidae | Lymantriinae | na | N | C | leaf | (Holloway 1999) |
| *Chrysocale gigantea* | Erebidae | Arctiinae | Ctenuchini | N | C | no | Expert J. Barbut |
| *Chrysocale regalis* | Erebidae | Arctiinae | Ctenuchini | N | C | no | Expert J. Barbut |
| *Cocytia durvillii* | Erebidae | Erebinae | Cocytiini | D | O | no | (Holloway et al. 2001) |
| *Corematura aliaria* | Erebidae | Arctiinae | Ctenuchini | D | O | b/w | (Project Noah 2017) |
| *Cosmosoma myrodora* | Erebidae | Arctiinae | Ctenuchini | N | O | b/w | (Ferro and Teston 2009) |
| *Dysschema boisduvalii* | Erebidae | Arctiinae | Pericopini | N | C | no | Expert J. Barbut |
| *Dysschema sacrifica* | Erebidae | Arctiinae | Pericopini | N | C | no | Expert J. Barbut |
| *Hyaleucerea vulnerata* | Erebidae | Arctiinae | Ctenuchini | N | C | b/w | (Weller et al. 2000) |
| *Hyalurga fenestra* | Erebidae | Arctiinae | Pericopini | D | O | no | (Poole 1970) |
| *Hypercompe robusta* | Erebidae | Arctiinae | Arctiini | N | O | no | (Hall 2014) |
| *Pantana sinica* | Erebidae | Lymantriinae | na | N | C | no | (Sastry et al. 1988) |
| *Phragmatobia fuliginosa* | Erebidae | Arctiinae | Spilosomini | N | O | no | (Crabo et al. 2020) |
| *Praeamastus fulvizonata* | Erebidae | Arctiinae | Arctiini | N | O | b/w | (Rab Green et al. 2011) |
| *Pseudohemihyalea rhoda* | Erebidae | Arctiinae | Arctiini | N | C | no | Expert J. Barbut |
| *Senecauxia coraliae* | Erebidae | Arctiinae | Phaegopterini | N | C | no | Expert J. Barbut |
| *Trichura mathina* | Erebidae | Arctiinae | Ctenuchini | D | O | b/w | Expert J. Barbut |
| *Hibrildes venosa* | Eupterotidae | Janinae | na | N | O | no | (Ansorge 1899) |
| *Nagara vitrea* | Euteliidae | Stictopterinae | na | N | C | no | (Comisión Nacional para la Gestión de la Biodiversidad (Costa Rica) 2018) |
| *Krananda extranotata* | Geometridae | Ennominae | na | N | C | leaf | (Arandhara and Tariang 2018) |
| *Pseudasellodes fenestraria* | Geometridae | Sterrhinae | na | N | O | leaf | (Brehm 2002) |
| *Trocherateina pohliata* | Geometridae | Larentiinae | na | D | O | no | Expert J. Barbut |
| *Zamarada sp* | Geometridae | Ennominae | na | N | O | leaf | Expert J. Barbut |
| *Macrosoma albipannosa* | Hedylidae | na | na | N | C | no | Expert J. Barbut |
| *Macrosoma conifera* | Hedylidae | na | na | N | C | no | Expert J. Barbut |
| *Epargyreus ridens* | Hesperiidae | na | na | D | O | no | (Shapiro 1978) |
| *Jemadia gnetus* | Hesperiidae | na | na | D | O | no | (Orellana 2010) |
| *Oxynetra semihyalina* | Hesperiidae | na | na | D | O | no | (Warren and Grishin 2017) |
| *Phanus vitreus* | Hesperiidae | na | na | D | O | no | (Corro-Chang 2017) |
| *Heterogynis penella* | Heterogynidae | na | na | D | O | no | (de Freina 2015) |
| *Semioptila fulveolans* | Himantopteridae | na | na | D | C | no | Expert J. Barbut |
| *Gastropacha africana* | Lasiocampidae | Lasiocampinae | na | N | O | leaf | (Muséum national d’Histoire naturelle 2003*a*) |
| *Pseudopsyche dembowskii* | Limacodidae | Limacodinae | na | N | C | no | Expert J. Barbut |
| *Megalopyge sp* | Megalopygidae | na | na | N | C | b/w | (Avilán et al. 2010) |
| *Gaujonia arbosi* | Noctuidae | Pantheinae | na | N | C | no | Expert J. Barbut |
| *Acraea braesia* | Nymphalidae | Heliconiinae | Acraeini | D | O | no | Expert J.Pierre |
| *Acraea iturina* | Nymphalidae | Heliconiinae | Acraeini | D | O | no | Expert J.Pierre |
| *Acraea necoda* | Nymphalidae | Heliconiinae | Acraeini | D | O | no | Expert J.Pierre |
| *Acraea pentapolis* | Nymphalidae | Heliconiinae | Acraeini | D | O | no | Expert J.Pierre |
| *Acraea semivitrea* | Nymphalidae | Heliconiinae | Acraeini | D | O | no | Expert J.Pierre |
| *Amauris hyalites* | Nymphalidae | Danainae | Danaini | D | O | no | Expert R. Rougerie |
| *Athesis clearista* | Nymphalidae | Danainae | Ithomiini | D | C | no | Expert M.Elias |
| *Cythaerias aurorina* | Nymphalidae | Satyrinae | Haeterini | D | C | no | Expert J. Barbut |
| *Dircenna klugii* | Nymphalidae | Danainae | Ithomiini | D | C | no | (Douglas et al. 2007) |
| *Dulcedo polita* | Nymphalidae | Satyrinae | Haeterini | D | C | no | (Constantino 1995) |
| *Episcada hemixanthe* | Nymphalidae | Danainae | Ithomiini | D | C | no | (DeVries 1987) |
| *Eutresis hypereia* | Nymphalidae | Danainae | Ithomiini | D | C | no | Experts M.Elias/ R. Rougerie |
| *Greta morgane* | Nymphalidae | Danainae | Ithomiini | D | C | no | Expert M.Elias |
| *Haetera piera* | Nymphalidae | Satyrinae | Haeterini | D | C | no | Expert R. Rougerie |
| *Ideopsis vitrea* | Nymphalidae | Danainae | Danaini | D | C | no | (Gerold and Fremerey 2004) |
| *Ithomia xenos* | Nymphalidae | Danainae | Ithomiini | D | C | no | Expert M.Elias |
| *Lycorea ilione* | Nymphalidae | Danainae | Danaini | D | O | no | (Orlandin et al. 2019) |
| *Methona confusa* | Nymphalidae | Danainae | Ithomiini | D | C | no | (Willmott et al. 2017) |
| *Napeogenes sylphis* | Nymphalidae | Danainae | Ithomiini | D | C | no | (Willmott et al. 2017) |
| *Oleria makrena* | Nymphalidae | Danainae | Ithomiini | D | C | no | (Poole 1970) |
| *Parantica sita* | Nymphalidae | Danainae | Danaini | D | C | no | (Cheng et al. 2015) |
| *Pseudohaetera hypaesia* | Nymphalidae | Satyrinae | Haeterini | D | C | no | (Álvarez 1993) |
| *Thyridia psidii* | Nymphalidae | Danainae | Ithomiini | D | C | no | Expert M.Elias/J. Barbut |
| *Archon apollinus* | Papilionidae | Parnassiinae | na | D | O | no | Expert R. Rougerie |
| *Cressida cressida* | Papilionidae | Papilioninae | na | D | O | no | (Braby 2016) |
| *Lamproptera meges* | Papilionidae | Papilioninae | Leptocircini | D | O | no | (Hu et al. 2014) |
| *Parides hahneli* | Papilionidae | Papilioninae | na | D | O | no | (Grice et al. 2019) |
| *Parnassius glacialis* | Papilionidae | Parnassiinae | Parnassini | D | C | no | (Yagi et al. 2001) |
| *Parnassius mnemosyne* | Papilionidae | Parnassiinae | Parnassini | D | O | no | (European Environment Agency 2019) |
| *Aporia crataegi* | Pieridae | Pierinae | Pierini | D | O | no | Expert R. Rougerie |
| *Dismorphia orise* | Pieridae | Dismorphinae | na | D | C | no | (Hoskins n.d.) |
| *Dismorphia theucharila* | Pieridae | Dismorphinae | na | D | C | no | Expert R. Rougerie |
| *Acanthopsyche sierricola* | Psychidae | Oiketicinae | na | D | O | no | (Bourgogne 1986) |
| *Chalioides ferevitrea* | Psychidae | Oiketicinae | Acanthopsychini | N | O | no | (Chakravarthy and Sridhara 2016) |
| *Oreopsyche albida* | Psychidae | Oiketicinae | Oreopsychini | N | O | no | Expert J. Barbut |
| *Thyridopteryx ephemeraeformis* | Psychidae | Oiketicinae | na | N | O | no | Expert J. Barbut |
| *Tephris eulella* | Pyralidae | Phycitinae | na | N | O | no | Expert J. Barbut |
| *Chorinea sylphina* | Riodinidae | Riodininae | Riodinini | D | O | no | Expert J. Barbut |
| *Ithomeis astrea* | Riodinidae | Riodininae | Riodinini | D | O | no | (DeVries 1987) |
| *Ithomiola cascella* | Riodinidae | Riodininae | Riodinini | D | O | no | (Salazar et al. 2010) |
| *Styx infernalis* | Riodinidae | Euselasiinae | Stygini | D | C | no | Expert R. Rougerie/J. Barbut |
| *Attacus atlas* | Saturniidae | Saturniinae | Attacini | N | C | no | Expert R. Rougerie |
| *Heliconisa pagenstecheri* | Saturniidae | Hemileucinae | na | D | O | no | Expert R. Rougerie, (Seitz 1906) |
| *Holocerina angulata* | Saturniidae | Saturniinae | Micragonini | N | C | leaf | Expert R. Rougerie |
| *Neorcarnegia basirei* | Saturniidae | Ceratocampinae | na | N | O | leaf | (Blest 1957; Drechsel and Lampe 1996) |
| *Orthogonioptilum violascens* | Saturniidae | Ludiinae | na | N | C | leaf | (Rougeot 1959) |
| *Rothschildia erycina* | Saturniidae | Saturniinae | Attacini | N | C | no | (Janzen 1984; Hernández-Chavarría et al. 2004) |
| *Melittia indica* | Sesiidae | Sesiinae | Melittiini | D | O | b/w | (Gorbunov and Arita 1996) |
| *Sesia apiformis* | Sesiidae | Sesiinae | Sesiini | D | O | b/w | (Skinner 2009) |
| *Synanthedon hector* | Sesiidae | Sesiinae | Synanthedonini | D | O | b/w | Expert J. Barbut |
| *Cephonodes hylas* | Sphingidae | Macroglossinae | Dilophonotini | D | O | b/w | (Pittaway and Kitching 2000*a*) |
| *Hemaris fuciformis* | Sphingidae | Macroglossinae | Dilophonotini | D | O | b/w | (Muséum national d’Histoire naturelle 2003*b*) |
| *Hemaris radians* | Sphingidae | Macroglossinae | Dilophonotini | D | O | b/w | (Pittaway and Kitching 2000*b*) |
| *Dysodia speculifera* | Thyrididae | Thyridinae | na | N | C | leaf | Expert J. Barbut |
| *Thyris fenestrella* | Thyrididae | Thyridinae | na | D | O | leaf | (Muséum national d’Histoire naturelle 2003*b*) |
| *Illiberis tenuis* | Zygaenidae | Procridinae | na | D | O | no | (Tarmann 2005) |
| *Phacusa translucida* | Zygaenidae | Procridinae | na | N | O | no | Expert J. Barbut |
| *Pryeria sinica* | Zygaenidae | Zygaeninae | na | D | O | no | (Yazaki et al. 2019) |

Species information included daytime activity (diurnal(D)/nocturnal(N)), habitat openness (open (O)/closed(C)) and mimicry (bee/wasp (b/w), leaf or non-mimic (no)). Ecological information was gathered by consulting experts, peer-reviewed papers and websites sponsored by museum collections or universities.

**Additional Material and Methods**

The rather cryptic butterfly colour was chosen based on 120 reflectance measures of evergreen oak *Quercus ilex* trunks. Reflectance measurements were taken using a spectrophotometer (Starline Avaspec-2048 L, Avantes) and a deuterium halogen lamp (Avalight DHS, Avantes) emitting in the 300-700 nm range, including UV, to which some predators of butterfly and moths, such as birds, are sensitive (Chen and Goldsmith 1986). Measurements were done relative to a white reference (lights turned on with no sample) and a dark reference (light turned off with no sample). For reflectance measurements, we used an optic probe (FC-UV200-2-1.5 x 100, Avantes) merging illumination and collection angles. Grey wings (Grey155: R=155, G=155, B=155) were printed on sketch paper Canson© using a HP Officejet Pro 6230 printer. Paper wings were laminated with a Polyester Opale Mat 75µm poach. We calculated colour and brightness contrast between oak trunk and artificial moth coloration, applying Vorobyev and Osorio discriminability model (Vorobyev and Osorio 1998) using *pavo* package with R software (Maia et al. 2013), UVS- (blue tit, (Hart et al. 2000)) and VS- bird vision (shearwater, (Hart 2001)) and forest shade as environment light (Gomez and Théry 2007). We found that artificial moths were chromatically similar (chromatic contrast of 0.47±0.16 JND for UVS vision and of 0.41±0.14 JND for VS vision), but lighter than oak trunks (achromatic contrast of 1.65±0.69 JND for UVS vision and of 1.65±0.68 JND for VS vision). For the butterflies that included transparent elements, two triangular windows of 252mm² each (56% of total artificial moth surface) were cut down from the grey triangle, and the remaining part was glued on top of one or six layers of transparent film 3M for inkjet printing. This transparent film was chosen as it is highly transparent, even in the UV range of the spectrum, and going from 1 to 6 transparent layers permitted a reduction of 50% of the transmittance between treatments (Fig. S2). Such transparent layer was coated with a transparent matte varnish to reduce its shininess, making them more similar to real Lepidoptera transparent wings that often exhibit nano antireflective structures (Pinna et al. 2020; Pomerantz et al. 2021). For transparency measurements, we measured specular transmittance from 300 to 1100nm, using a deuterium-halogen lamp (Avalight DHS, Avantes), optical fibres (FC-UV200-2-1.5 x 100, Avantes) and a spectrometer (Starline Avaspec-2048 L, Avantes). Fibres were separated, aligned and 5mm apart and the wing sample was placed perpendicular between them at equal distance (light spot of 1mm diameter). Spectra were taken relative to a dark (light off) and to a white reference (no sample between the fibres) measurement. On top of butterfly wings, we added an artificial grey body. These butterfly bodies were prepared with flour (428 g), lard (250 g) and water (36 g), following Carrol & Sherratt (2013). Yellow, red and blue food edible colourings were used to dye the pastry in grey, imitating artificial wing colour. Such malleable mix permits to register and distinguish marks made by bird beaks from insect jaws. Both paper wings and artificial bodies were measured in spectrometry, similarly to how trunk reflectance was measured (Fig. S2). Bodies could attract predator attention, but it has been previously shown that wingless bodies had the same attractiveness as that of butterflies with large transparent elements (Arias et al. 2020). Artificial butterflies and bodies were pinned to evergreen oak *Quercus ilex* trunks. To avoid ant attacks, Vaseline and a piece of highly sticky double-faced transparent tape were stuck between the wings and the trunk.

Table S1. Variation of mean proportion of transmitted light according to mimicry, habitat, daytime activity, wing size, wing and proportion of clearwing surface excluding (on the left) or including interactions between them (on the right). For Bayesian estimations see Tables S2 a, b, c and d.

|  |  | Model with single factors only | | | | Model with single factors and interactions | | | |
| --- | --- | --- | --- | --- | --- | --- | --- | --- | --- |
|  |  | Estimate ± se | DF | *t* |  | Estimate ± se | DF | *t* |  |
|  | **(Intercept)** | **58.15±3.1** | **502** | **18.77** | ******* | **61.7±3.56** | **488** | **17.32** | *** |
|  | ActHab (DayOpen>DayClosed) | 3.33±3.64 | 101 | 0.92 |  | 2.66±5.85 | 101 | 0.45 |  |
|  | **ActHab (Day>Night)** | 1.99±1.91 | 101 | 1.04 |  | **7.93±3.06** | **101** | **2.59** | ** |
|  | **ActHab (Open>Closed)** | -4.55±2.66 | 101 | -1.71 | ~ | **-13.16±4.13** | **101** | **-3.19** | ******* |
|  | **% clearwing** | **0.26±0.02** | **502** | **11.08** | ******* | 0.07±0.04 | 488 | 1.84 | ~ |
|  | **Mimicry:bee/wasp>leaf+non-mimics** | **4.91±1.91** | **101** | **2.58** | ****** | **9.45±2.25** | **101** | **4.19** | ******* |
|  | Mimicry:leaf<non-mimics | -5.02±2.96 | 101 | -1.7 | ~ | -4.1±3.75 | 101 | -1.1 |  |
|  | **Wing length** | **-0.43±0.07** | **502** | **-6.5** | ******* | **-0.38±0.1** | **488** | **-3.89** | ******* |
|  | **Wing (Forewing>Hindwing)** | **3.36±0.38** | **502** | **8.75** | ******* | **1.39±0.59** | **488** | **2.33** | ***** |
|  | **ActHab (DO>DC): % clearwing** |  |  |  |  | **0.16±0.05** | **488** | **2.96** | ****** |
|  | ActHab (D>N): % clearwing |  |  |  |  | 0.03±0.03 | 488 | 1.23 |  |
|  | ActHab (O>C): % clearwing |  |  |  |  | 0.07±0.04 | 488 | 1.58 |  |
|  | Mimicry (bw>l+n.m): % clearwing |  |  |  |  | 0.04±0.02 | 488 | 1.48 |  |
|  | **Mimicry (leaf<n.m): % clearwing** |  |  |  |  | **0.19±0.05** | **488** | **3.87** | ******* |
|  | **Mimicry (bw>l+n.m):W.length** |  |  |  |  | **-0.28±0.04** | **488** | **-6.18** | ******* |
|  | Mimicry (leaf<n.m):W.length |  |  |  |  | -0.16±0.09 | 488 | -1.88 | ~ |
|  | **ActHab (DO>DC):Wing length** |  |  |  |  | **-0.42±0.16** | **488** | **-2.63** | ****** |
|  | **ActHab (D>N):Wing length** |  |  |  |  | **-0.32±0.09** | **488** | **-3.78** | ******* |
|  | **ActHab (O>C):Wing length** |  |  |  |  | **0.38±0.14** | **488** | **2.81** | ****** |
|  | **ActHab (DO>DC):Wing (F>H)** |  |  |  |  | **2.2±0.86** | **488** | **2.57** | ****** |
|  | **ActHab (D>N):Wing (F>H)** |  |  |  |  | **1.92±0.45** | **488** | **4.28** | ******* |
|  | ActHab (O>C):Wing (F>H) |  |  |  |  | -1.16±0.68 | 488 | -1.7 | ~ |
|  | **Wing length:Wing (F>H)** |  |  |  |  | **0.05±0.02** | **488** | **2.12** | ***** |

Linear mixed model with transmittance as dependent variable and the following explanatory variables: the combination between daytime activity and habitat type (ActHab: nocturnal-open/nocturnal-closed/diurnal-open/diurnal-closed), wing length in mm, proportion of clearwing area (% clearwing), mimicry syndrome (bee/wasp -, leaf - or non-mimics), wing (fore-/hindwing) and the interactions: mimicry syndrome and proportion of clearwing area, mimicry syndrome and wing size, proportion of clearwing area and habitat type, wing size and habitat type, wing size and proportion of clearing area, wing and wing length and proportion of clearwing area, wing length and habitat as explanatory variables. Wing measurements nested in species was considered as random effect. LMM p values below 0.05 are statistically significant. *** stands for p<0.001, ** for p<0.01, * for p<0.05 and ~ for p<0.1.

Table S2. Bayesian phylogenetic models using mean percentage of transmitted light as dependant variable excluding (Tables S2a-d, left part) or including interactions between them (Tables S2a-d, right part). Specific mimicry syndrome (bee/wasp mimics vs. all other species, and leaf mimics vs. non-mimics) and daytime activity/habitat (ActHab: diurnal vs. nocturnal, open vs. closed habitats, and diurnal open habitat vs. diurnal closed habitat) contrast were tested in the different models. Fitted models additionally include wing length in mm, proportion of clearwing area (% clearwing) and wing (fore-/hindwing) as explanatory variables. Bayesian estimates whose confidence interval did not include 0 are statistically significant. The summary table below shows which levels were tested in which models.

|  | 2A | 2B | 2C | 2D |
| --- | --- | --- | --- | --- |
| **(Intercept)** | **x** | **x** |  |  |
| ActHab (DayOpen>DayClosed) |  |  | x |  |
| ActHab (Day>Night) | x |  |  |  |
| **ActHab (Open>Closed)** |  | x |  |  |
| **% clearwing surface** | **x** | **x** |  |  |
| **Mimicry:bee/wasp>leaf+non-mimics** | **x** | **x** |  |  |
| Mimicry:leaf<non-mimics |  |  |  | x |
| **Wing length** | **x** | **x** |  |  |
| **Wing (Forewing>Hindwing)** | **x** | **x** |  |  |
| **ActHab (DO>DC): % clearwing** |  |  | **x** |  |
| **ActHab (D>N): % clearwing** | **x** |  |  |  |
| **ActHab (O>C): % clearwing** |  | **x** |  |  |
| Mimicry (bw>l+n.m): % clearwing | x | x |  |  |
| **Mimicry (leaf<n.m): % clearwing** |  |  |  | **x** |
| **Mimicry (bw>l+n.m):W.length** | **x** | **x** |  |  |
| Mimicry (leaf<n.m):W.length |  |  |  | x |
| ActHab (DO>DC):Wing length |  |  | x |  |
| **ActHab (D>N):Wing length** | x |  |  |  |
| ActHab (O>C):Wing length |  | x |  |  |
| **ActHab (DO>DC):Wing (F>H)** |  |  | **x** |  |
| **ActHab (D>N):Wing (F>H)** | **x** |  |  |  |
| **ActHab (O>C):Wing (F>H)** |  | **x** |  |  |
| **Wing length:Wing (F>H)** |  | **x** |  |  |

Table S2a. Model testing the contrast mimicry bee/wasp mimics > leaf+no mimics and ActHab: diurnal > nocturnal. Including all data

|  | Without interaction | With interaction |
| --- | --- | --- |
|  | Estimate [CI95%] | Estimate [CI95%] |
| **(Intercept)** | **49.26[22.88,75.66]** | **52.2[25.77,79]** |
| ActHab (Diurnal>Nocturnal) | 0.46[-10.07,10.82] | -5.2[-18.52,7.72] |
| **% clearwing surface** | **0.26[0.21,0.31]** | **0.3[0.23,0.36]** |
| **Mimicry:bw>l+n.m** | **26.38[15.16,37.12]** | **36.97[23.3,50.28]** |
| **Wing length** | **-0.5[-0.63,-0.37]** | **-0.56[-0.72,-0.41]** |
| **Wing (F>H)** | **-8.01[-9.54,-6.43]** | **-10[-13.15,-6.93]** |
| **ActHab (D>N): % clearwing surf.** |  | **-0.18[-0.28,-0.08]** |
| Mimicry (bw>l+n.m): % clearwing surf. |  | 0.06[-0.09,0.2] |
| **Mimicry (bw> l+n.m):W.length** |  | **-0.73[-1.04,-0.41]** |
| **ActHab (D>N):Wing length** |  | **0.46[0.13,0.77]** |
| **ActHab (D>N):Wing (F>H)** |  | **8.41[4.76,12.12]** |
| Wing length:Wing (F>H) |  | -0.04[-0.14,0.05] |
| **phylogenetic.variance** | **883.51[650.87,1172.46]** | **851.34[630.54,1132.73]** |
| **residual.variance** | **73.47[67.18,80.39]** | **69.41[63.51,76.13]** |

Table S2b. Model testing the contrast mimicry: bee/wasp mimics > leaf+no mimics and ActHab: open> closed. Including all data

|  | Without interaction | With interaction |
| --- | --- | --- |
|  | Estimate [CI95%] | Estimate [CI95%] |
| **(Intercept)** | **51.89[26.09,78.35]** | **57.21[28.01,85.76]** |
| **ActHab (Open>Closed)** | -3.67[-11.9,4.66] | **-16.98[-30.04,-4.19]** |
| **% clearwing surface** | **0.26[0.21,0.3]** | 0.05[-0.02,0.12] |
| **Mimicry:bw>l+n.m** | **27.21[16.25,38.12]** | **41.89[28.25,55.61]** |
| **Wing length** | **-0.5[-0.63,-0.36]** | **-0.5[-0.72,-0.29]** |
| **Wing (F>H)** | **-7.99[-9.58,-6.42]** | -2.6[-5.61,0.41] |
| **ActHab (O>C): % clearwing surf.** |  | **0.38[0.28,0.48]** |
| Mimicry (bw>l+n.m): % clearwing surf. |  | -0.07[-0.2,0.07] |
| **Mimicry (bw> l+n.m):W.length** |  | **-0.67[-0.98,-0.37]** |
| ActHab (O>C):Wing length |  | 0[-0.26,0.27] |
| **ActHab (O>C):Wing (F>H)** |  | **-4.22[-7.32,-1.21]** |
| **Wing length:Wing (F>H)** |  | **-0.16[-0.25,-0.07]** |
| **phylogenetic.variance** | **879.63[661.88,1173.38]** | **948.28[709.43,1266.47]** |
| **residual.variance** | **73.59[67.09,80.49]** | **65.51[59.92,71.82]** |

Table S2c. Model testing the contrast ActHab: diurnal open> diurnal closed. Including only a subset of data that excluded nocturnal species.

|  | Without interaction | With interaction |
| --- | --- | --- |
|  | Estimate [CI95%] | Estimate [CI95%] |
| **(Intercept)** | **54.64[28.95,78.49]** | **67.06[38.41,94.5]** |
| ActHab (DayOpen>DayClosed) | 1.64[-8.88,12.88] | -17.24[-37.39,2.68] |
| **% clearwing surface** | **0.31[0.24,0.37]** | -0.01[-0.12,0.1] |
| **Mimicry:bw>l+n.m** | 12.19[-4.64,29.32] | **26.01[6.38,46.45]** |
| **Wing length** | **-0.6[-0.76,-0.44]** | **-0.58[-0.89,-0.28]** |
| **Wing (F>H)** | **-10.28[-12.25,-8.34]** | -1.53[-7.09,4.04] |
| **ActHab (DO>DC): % clearwing surf.** |  | **0.44[0.3,0.58]** |
| Mimicry (bw>l+n.m): % clearwing surf. |  | -0.01[-0.18,0.16] |
| **Mimicry (bw> l+n.m):W.length** |  | **-0.98[-1.4,-0.55]** |
| ActHab (DO>DC):Wing length |  | 0.06[-0.3,0.41] |
| **ActHab (DO>DC):Wing (F>H)** |  | **-5.83[-10.07,-1.6]** |
| **Wing length:Wing (F>H)** |  | **-0.23[-0.4,-0.07]** |
| **phylogenetic.variance** | **560.88[375.58,828.14]** | **598.97[400.55,879.85]** |
| **residual.variance** | **81.72[72.52,92.04]** | **70.05[62.13,78.58]** |

Table S2d. Model testing the contrast mimicry: leaf < other non-mimics clearwing species. Including only a subset of data that excluded bee/wasps mimics.

|  | Without interaction | With interaction |
| --- | --- | --- |
|  | Estimate [CI95%] | Estimate [CI95%] |
| **(Intercept)** | **52.29[25.5,80.08]** | **53.98[25.87,82]** |
| **ActHab (Open>Closed)** | -2.7[-11.88,6.34] | **-16.02[-30.31,-2.53]** |
| **% clearwing surface** | **0.25[0.2,0.31]** | **0.07[0,0.14]** |
| Mimicry:l<n.m | 0.87[-13.12,15.31] | 5.09[-12.46,22.93] |
| **Wing length** | **-0.49[-0.63,-0.35]** | **-0.5[-0.73,-0.28]** |
| Wing (F>H) | **-8.94[-10.57,-7.25]** | -1.66[-4.78,1.46] |
| **ActHab (O>C): % clearwing surf.** |  | **0.4[0.3,0.51]** |
| **Mimicry (l<n.m): % clearwing surf.** |  | **-0.37[-0.57,-0.19]** |
| Mimicry (l<n.m):W.length |  | 0.3[-0.03,0.64] |
| ActHab (O>C):Wing length |  | 0.01[-0.27,0.29] |
| **ActHab (O>C):Wing (F>H)** |  | **-4.24[-7.53,-1.09]** |
| **Wing length:Wing (F>H)** |  | **-0.17[-0.27,-0.08]** |
| **phylogenetic.variance** | **883.3[644.68,1186.24]** | **958.43[694.16,1301.67]** |
| **residual.variance** | **77.66[70.24,85.4]** | **69.07[62.42,76.07]** |

Table S3. PGLS results for ActHab, wing length (rows 1-4) and proportion of clearwing surface (rows 5-8) for both forewing and hindwing.

|  |  | Forewing | | | | Hindwing | | |
| --- | --- | --- | --- | --- | --- | --- | --- | --- |
|  |  | Estimate± se | *t* | |  | Estimate± se | *t* |  |
| Wing length | **Intercept** | **28.71±11.16** | **2.57** | ****** | | **19.44±8.35** | **2.33** | ***** |
|  | ActHab.DO>DC | -1.13±2.74 | -0.41 |  | | -0.38±2.42 | -0.16 |  |
|  | ActHab.D>N | 1.49±1.41 | 1.06 |  | | 1.82±1.14 | 1.60 |  |
|  | **ActHab.O>C** | **-4.52±2.13** | **-2.12** | ***** | | -1.98±1.74 | -1.14 |  |
| Prop. of clearwing surface | **Intercept** | **46.1±18.53** | **2.49** | ****** | | **46.98±17.97** | **2.61** | ****** |
|  | **ActHab.DO>DC** | **-19.78±4.55** | **-4.34** | ******* | | **-10.59±5.21** | **-2.03** | ***** |
|  | ActHab.D>N | 3.12±2.34 | 1.33 |  | | 0.55±2.45 | 0.23 |  |
|  | **ActHab.O>C** | **14.19±3.53** | **4.02** | ******* | | **7.54±3.75** | **2.01** | ***** |

*** stands for statistical significance below 0.001, ** for p<0.01, * for p<0.05, ~for p<0.1.

Table S4. Results of coevolution tests between mimicry and scale characteristics

| Mimicry levels included | Wing | Scale characteristic | L. Dep | L. Indep | LRT | df | *p* |  |
| --- | --- | --- | --- | --- | --- | --- | --- | --- |
| Bee/wasp and leaf mimics | F | Presence | -30.58 | -30.79 | 0.42 | 4 | 0.98 |  |
|  |  | Scale type | -23.29 | -25.73 | 4.88 | 4 | 0.3 |  |
|  |  | Color | -27.29 | -28.41 | 2.24 | 4 | 0.69 |  |
|  |  | Insertion | -30.97 | -31.21 | 0.49 | 4 | 0.97 |  |
|  | H | **Presence** | **-29.05** | **-34.27** | **10.45** | **4** | **0.033** | ***** |
|  |  | Scale type | -22.3 | -23.61 | 2.62 | 4 | 0.62 |  |
|  |  | Color | -25.57 | -25.86 | 0.58 | 4 | 0.96 |  |
|  |  | **Insertion** | **-27.8** | **-33.33** | **11.06** | **4** | **0.026** | ***** |
| No- and bee/wasp mimics | F | Presence | -82.9 | -85.1 | 4.4 | 4 | 0.35 |  |
|  |  | Scale type | -91.09 | -91.12 | 0.06 | 4 | 1 |  |
|  |  | Color | -98.5 | -100.52 | 4.04 | 4 | 0.4 |  |
|  |  | Insertion | -100.82 | -100.91 | 0.18 | 4 | 1 |  |
|  | H | **Presence** | **-73.46** | **-88.46** | **29.99** | **4** | **0.001** | ****** |
|  |  | Scale type | -93.9 | -95.18 | 2.56 | 4 | 0.633 |  |
|  |  | **Color** | **-88.75** | **-99.43** | **21.34** | **4** | **0.001** | ****** |
|  |  | **Insertion** | **-95.49** | **-101.07** | **11.17** | **4** | **0.025** | ***** |
| No- and leaf mimics | F | Presence | -71.32 | -71.81 | 0.98 | 4 | 0.913 |  |
|  |  | Scale type | -74.59 | -77.31 | 5.45 | 4 | 0.244 |  |
|  |  | **Color** | **-84.5** | **-90.31** | **11.62** | **4** | **0.02** | ***** |
|  |  | Insertion | -90.51 | -90.63 | 0.24 | 4 | 0.993 |  |
|  | H | Presence | -70.57 | -71.12 | 1.09 | 4 | 0.86 |  |
|  |  | **Scale type** | **-84.27** | **-88.93** | **9.33** | **4** | **0.053** | ***** |
|  |  | **Color** | **-86.66** | **-93.1** | **12.87** | **4** | **0.012** | ***** |
|  |  | Insertion | -95.63 | -96.14 | 1.01 | 4 | 0.91 |  |

Likelihood of dependent (L. Dep. Coevolution, 4 parameters estimated) and independent (L. Indep, 8 parameters estimated) evolution between pairs of binary characters per wing (F: forewing and H: hindwing). As mimicry has three levels, subsets including two of them were used and reported in the first column. Scale traits included in the analyses were: presence/absence, type (lamellar scales, piliform scales or both), colour (transparent/coloured), and insertion (erect/flat). *** stands for statistical significance below 0.001, ** for p<0.01, * for p<0.05, ~for p<0.1.

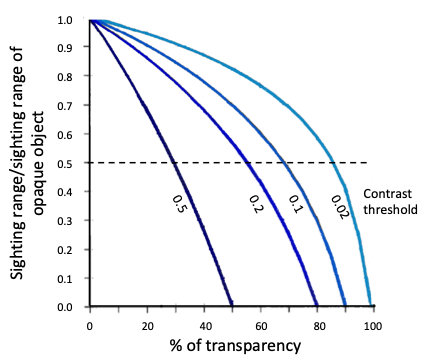

Figure S1. Sighting distance vs percentage of transparency of a prey (assumed to be independent of wavelength), according to the contrast threshold (lighter blue lines for lower contrast thresholds) of the visual system viewing the prey. transparent prey would be detected at twice the sighting range (horizontal dashed line, as an example) relative to the sighting range of a fully opaque prey for transparency higher than 90% for contrast threshold of 0.02, and higher than 30% for contrast threshold of 0.5. Figure adapted from Johnsen and Widder (1998). Contrast threshold is known to decrease as ambient light levels increase (Johnsen and Widder 1998) and light blue curves can be encountered for species living in shallower zones than dark blue curves.

a

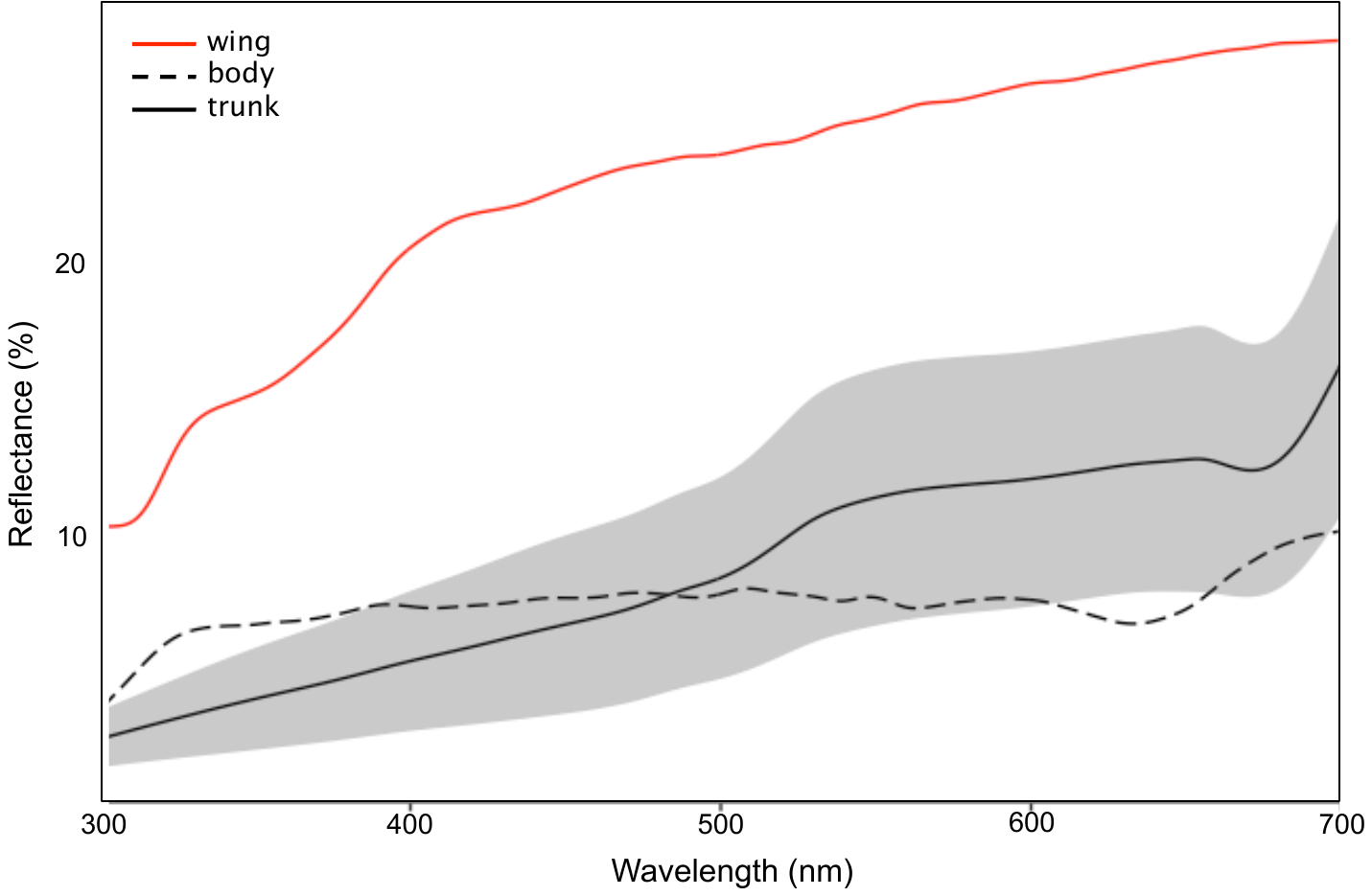

b

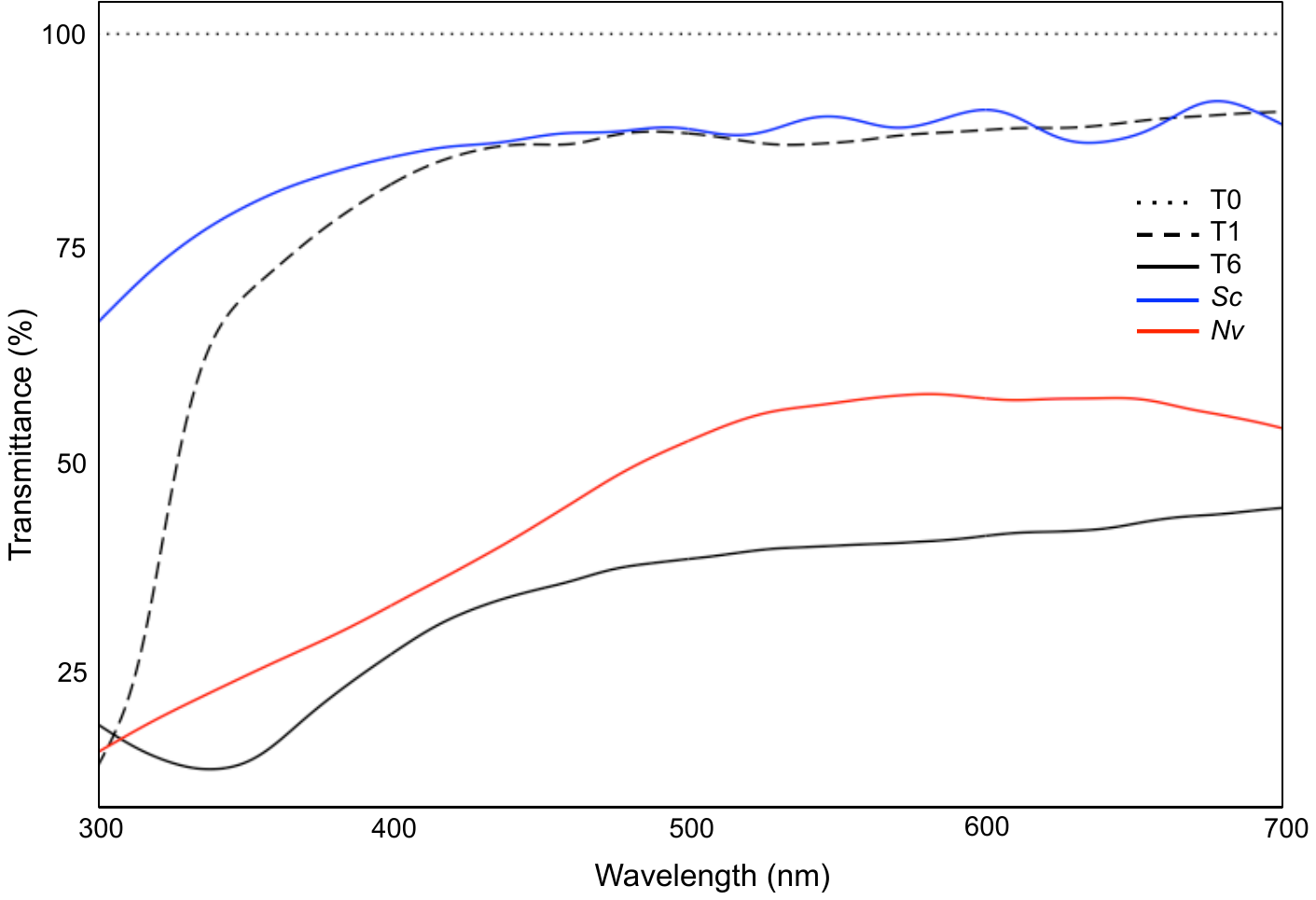

Figure S2. a) Reflectance of evergreen oak trunks (mean represented by solid line with a confidence interval of ± 1 standard deviation), artificial butterfly bodies (dashed line) and completely opaque butterfly “C” (red line). b) Percentage of transmittance of the different treatments used in the field experiment: poorly transparent butterfly with 6 layers of transparent film “T6”, highly transparent butterfly with a single layer of transparent film “T1”, and fully transparent butterfly with no film in the transparent zones “T0”. Transmittance measurements were done after coating the transparent film with the opaque varnish. Additionally, transmittance of two Lepidoptera species exhibiting transparency are illustrated: in *Senecauxia coraliae* (Sc, in blue), and in *Nagara vitrea* (Nv, in red)*.*

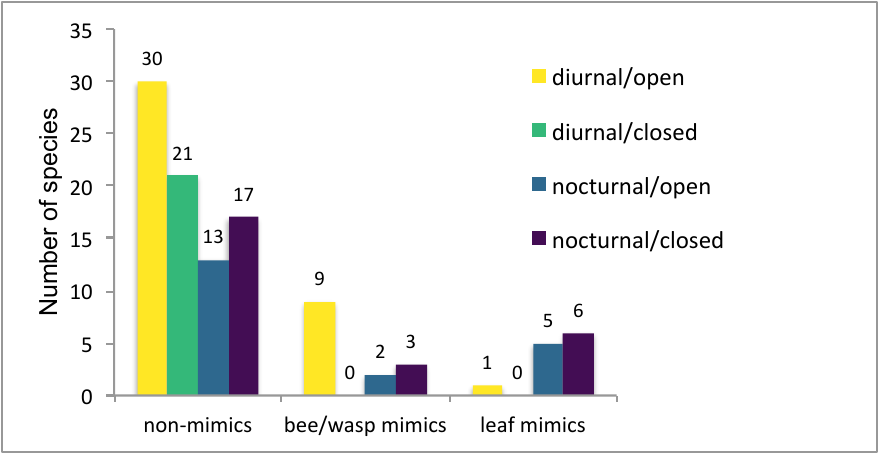

Figure S3. Distribution of species used on this study according to their daytime activity, habitat and mimicry.

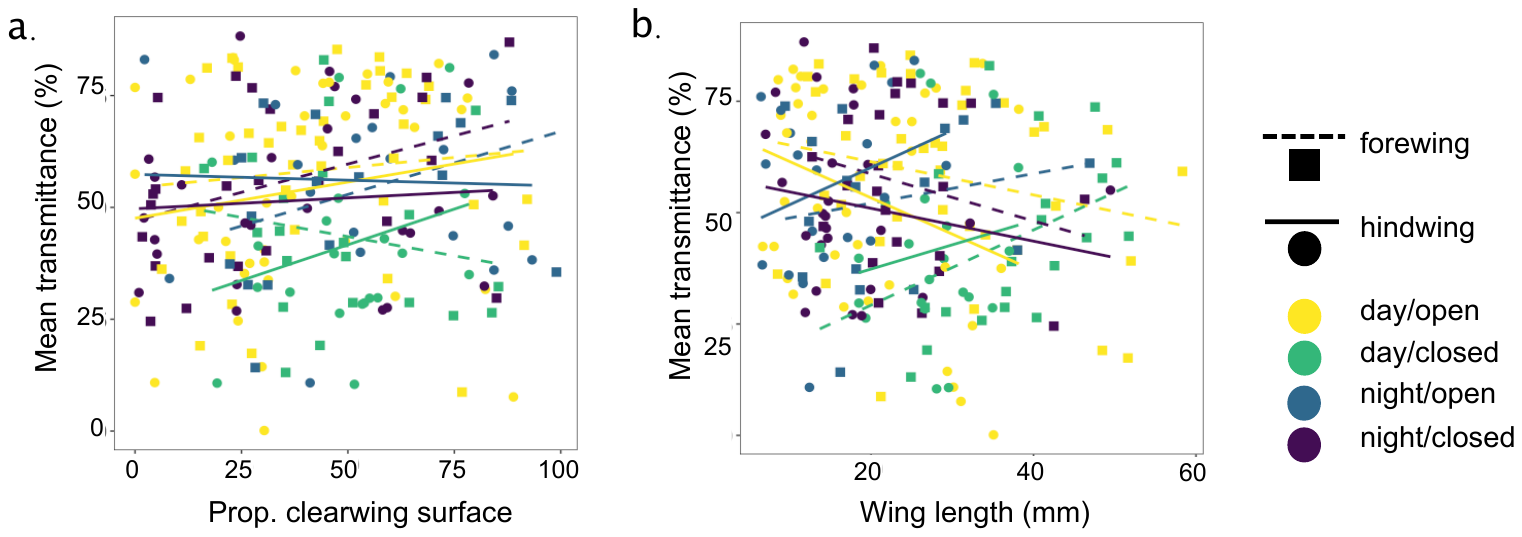

Figure S4: Relationship between mean transmittance for each ActHab category (combining daytime activity and habitat: diurnal/open (yellow), diurnal/closed (green), nocturnal/open (blue) and nocturnal/closed (purple)), wing (forewing: square/dashed and hindwing: circle/plain) and a) proportion of clearwing surface or b) wing length. Plotted lines in a. and b. correspond to linear regressions per wing and ActHab level. Values larger than 60 mm were not plotted for clarity reasons but were included in the analyses.

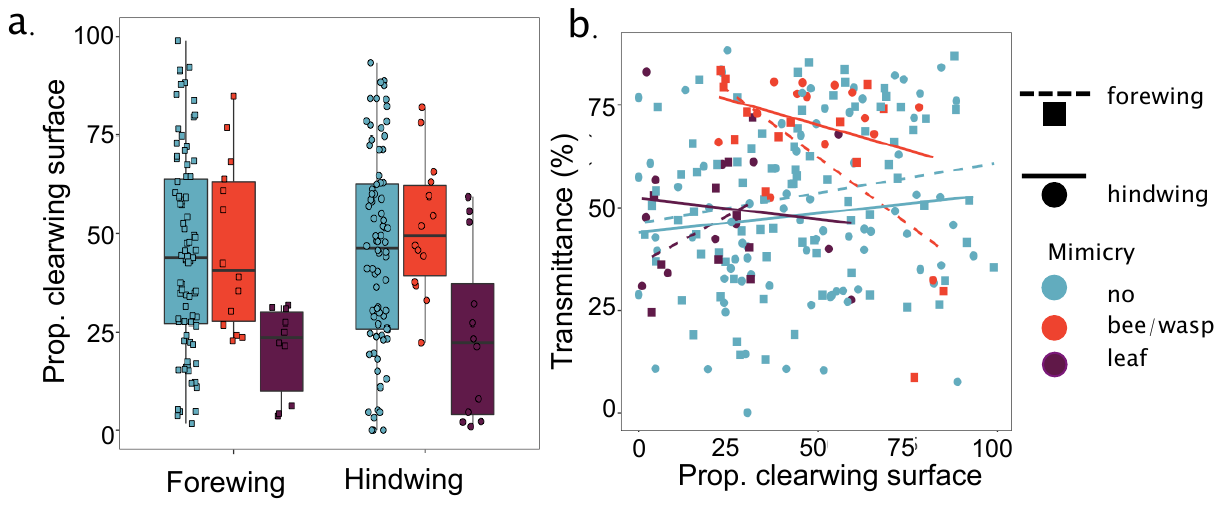

Figure S5: Variation between bee/wasp mimics (red), leaf mimics (purple) and no mimics (blue) in a) proportion of clearwing surface and b) in the relationship between wing size and mean light transmittance. Plotted lines in b. correspond to linear regressions per wing and mimicry level.

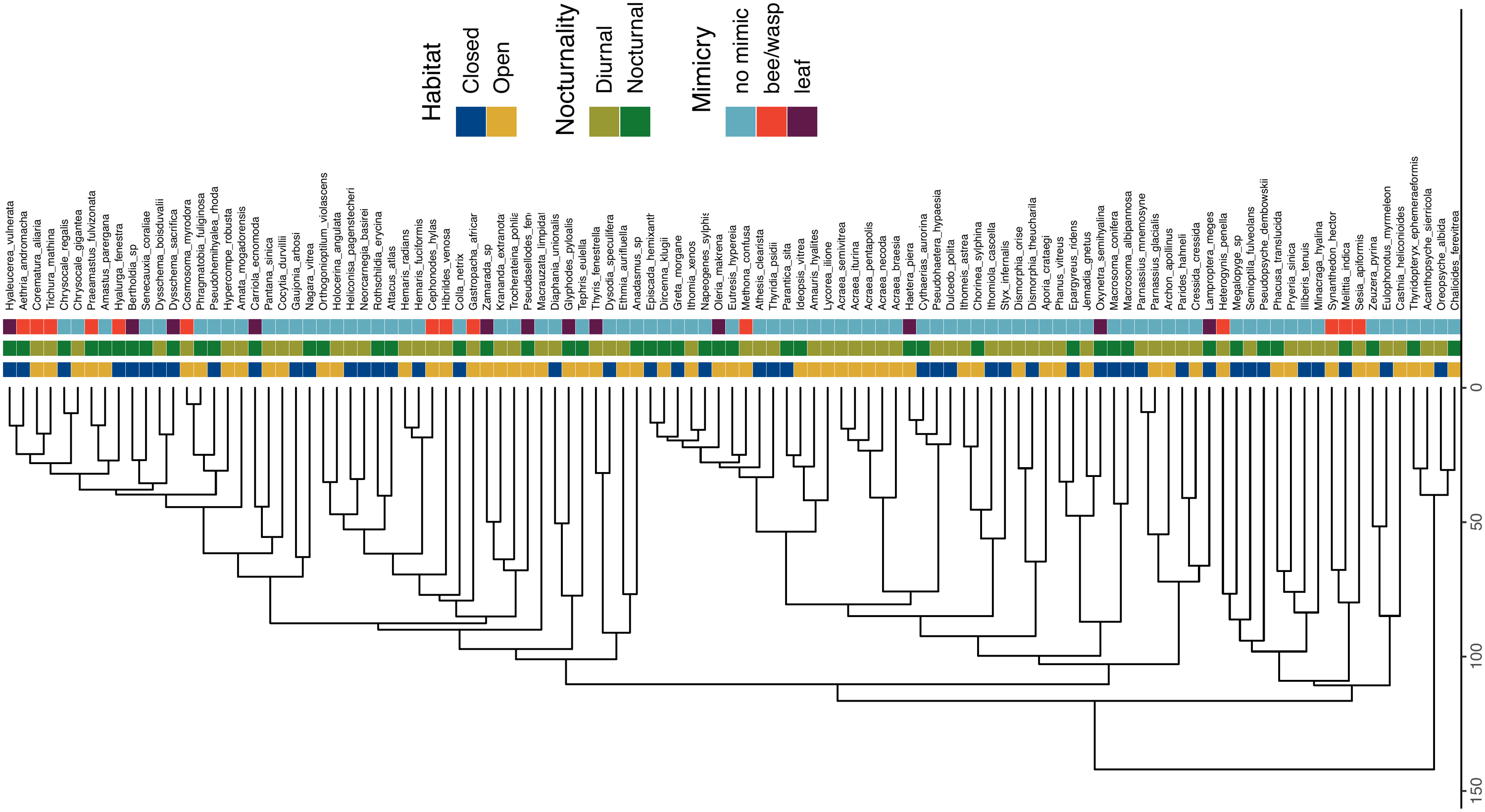

Figure S6. Phylogeny and daytime activity, habitat openness and mimicry for the 107 species of Lepidoptera

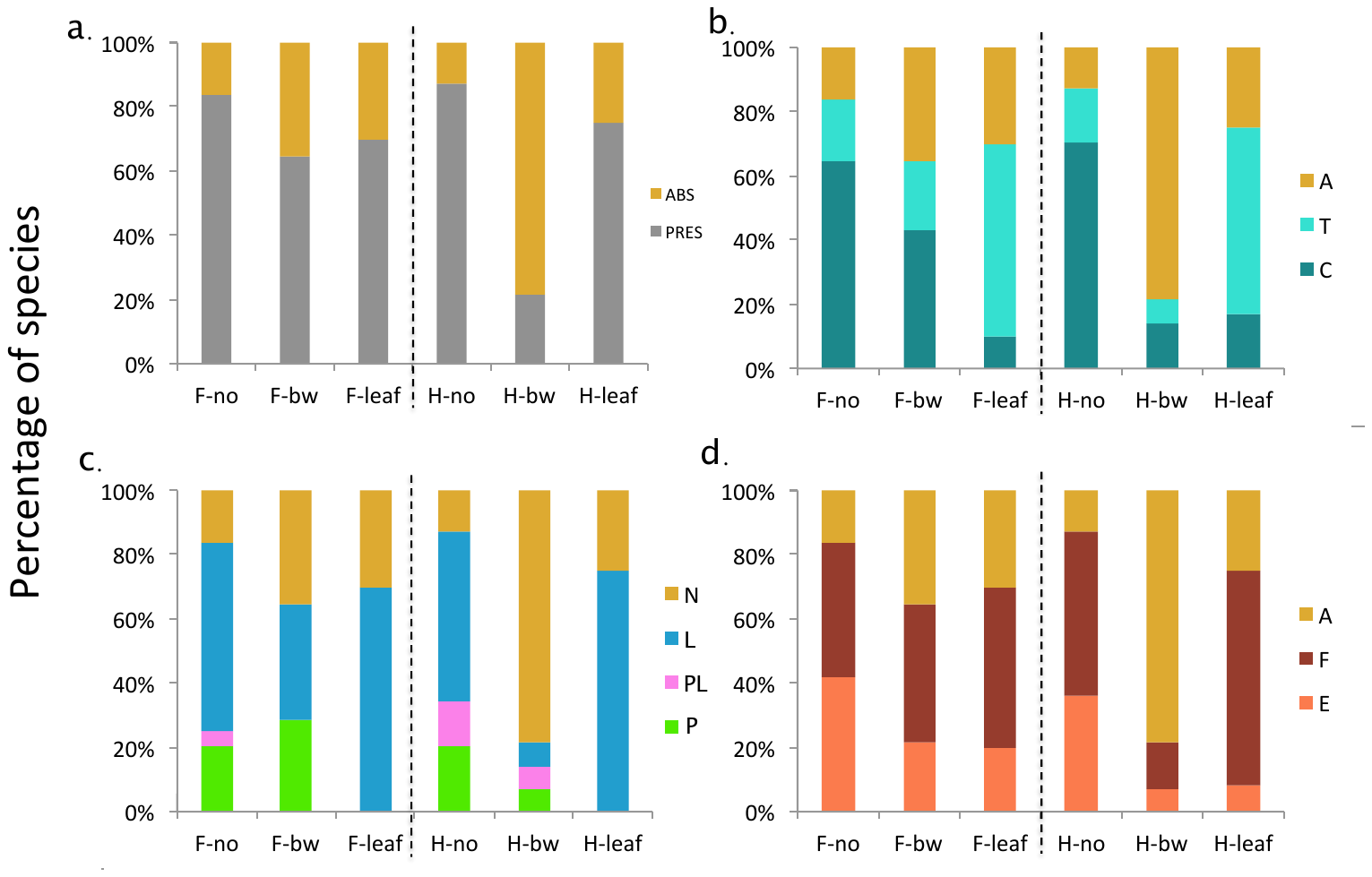

Figure S7. Distribution of a) presence/absence of scales, b) scales coloured (A: absent/ T: transparent/ C: coloured), c) scale type (N: no scales/ L: lamellar scales/PL: lamellar scale and piliform scales/ P: piliform scales) and d) scale insertion (A: absent/ F: flat/ E: erect) in no mimics (no), bee/wasp mimics (bw), and leaf mimics (leaf), in forewing (F) and hindwing (H).

References

Álvarez, J. 1993. Inventario de las mariposas (Lepidoptera: Rhopalocera), Con anotaciones ecológicas, para dos zonas del Departamento de Risaralda, Colombia. Trabajo de grado (Biología). Facultad de Ciencias, Universidad Nacional, Bogotá.

Ansorge, W. J. 1899. Under the African Sun: A Description of Native Races in Uganda, Sporting Adventures and Other Experiences. Longmans, Green and Co., London.

Arandhara, S., and R. R. Tariang. 2018. Drivers regulating species composition of the larger nocturnal moths in Tinsukia district, Assam. Journal of Entomology and Zoology Studies 6:748–55.

Arias, M., M. Elias, C. Andraud, S. Berthier, and D. Gomez. 2020. Transparency improves concealment in cryptically coloured moths. Journal of Evolutionary Biology 33:247–252.

Avilán, L., B. Guerrero, E. Álvarez, and A. Rodríguez-Acosta. 2010. Description of envenomation by the “gusano-pollo” caterpillar (Megalopyge opercularis) in Venezuela. Investigación clínica 51:127–132.

Beccacece, H. M. 2017. A new species of the genus Bertholdia Schaus, 1896 (Lepidoptera: Erebidae: Arctiinae) from the Neotropical region: Bertholdia zoenia sp. n. Zootaxa 4238:zootaxa-4238.

Blest, A. 1957. The evolution of protective displays in the Saturnioidea and Sphingidae (Lepidoptera). Behaviour 11:257–309.

Bourgogne, J. 1986. Une espèce peu connue: Acanthopsyche carbonaria Karsh [Lep. Psychidae]. Bulletin de la Société entomologique de France 91:287–291.

Braby, M. 2016. The complete field guide to butterflies of Australia. Csiro Publishing.

Brehm, G. 2002. *Diversity of geometrid moths in a montane rainforest in Ecuador*. Universität Bayreuth, Germany.

Carroll, J., and T. Sherratt. 2013. A direct comparison of the effectiveness of two anti‐predator strategies under field conditions. Journal of Zoology 291:279–285.

Chakravarthy, A. K., and S. Sridhara. 2016. Economic and ecological significance of arthropods in diversified ecosystems: Sustaining regulatory mechanisms. Springer.

Chen, D.-M., and T. H. Goldsmith. 1986. Four spectral classes of cone in the retinas of birds. Journal of Comparative Physiology A 159:473–479.

Cheng, W. W., H. S. Pun, O. Chung, T. Fukumura, I. Kanazawa, and F. Y. B. Reserve. 2015. Parantica sita niphonica (Lepidoptera: Nymphalidae) migrated from Japan to Hong Kong, southern China in 2013. Bulletin of the Osaka Museum of Natural History 69.

Comisión Nacional para la Gestión de la Biodiversidad (Costa Rica). 2018. Plataforma Informática para la gestión del conocimiento y la información nacional sobre Biodiversidad. Database.

Constantino, L. M. 1995. Revisión de la tribu Haeterini Herrich-Schäffer, 1864 en Colombia (Lepidoptera: Nymphalidae, Satyrinae). SHILAP Revista Lepidopterológica 23:49–76.

Corro-Chang, P. E. 2017. Behavioural notes and attraction on Lepidoptera around the Gehry’s Biodiversity Museum (Causeway, Calzada de Amador, Panamá, República de Panamá). Biodiversity data journal.

Crabo, L., R. Zack, and M. Peterson. 2020. Phragmatobia fuliginosa (Linnaeus, 1758). Pacific Northwest Moths.

de Freina, J. J. 2015. Beitrag zur Artengruppe von Heterogynis penella (Hübner, 1819 [„1816 “]) auf der Iberischen Halbinsel: Beschreibung von H. segurana sp. n., mit Ergänzungen zu anderen Arten (Lepidoptera: Zygaenoidea, Heterogynidae).

DeVries, P. J. 1987. The butterflies of Costa Rica and their natural history. Papilionidae, Pieridae, Nymphalidae. Princeton Univ. Press: xxii.

Douglas, J. M., T. W. Cronin, T.-H. Chiou, and N. J. Dominy. 2007. Light habitats and the role of polarized iridescence in the sensory ecology of neotropical nymphalid butterflies (Lepidoptera: Nymphalidae). Journal of Experimental Biology 210:788–799.

Drechsel, U., and R. E. Lampe. 1996. Die Präimaginalstadien von Neorcarnegia basirei (Schaus 1892) mit Anmerkungen zur Biogeographie (Lepidoptera: Saturniidae, Ceratocampinae). Nachrichten des Entomologischen Vereins Apollo, Frankfurt am Main, NF 17:143–152.

European Environment Agency. 2019. Clouded Apollo- Parnassius mnemosyne (Linnaeus, 1758).

Ferro, V. G., and J. A. Teston. 2009. Composição de espécies de Arctiidae (Lepidoptera) no sul do Brasil: relação entre tipos de vegetação e entre a configuração espacial do hábitat. Revista Brasileira de entomologia 53:278–286.

Gerold, G., and M. Fremerey. 2004. Land use, nature conservation and the stability of rainforest margins in southeast Asia. Springer Science & Business Media.

Gomez, D., and M. Théry. 2007. Simultaneous crypsis and conspicuousness in color patterns: comparative analysis of a neotropical rainforest bird community. the american naturalist 169:S42–S61.

Gorbunov, O. G., and Y. Arita. 1996. New and little-known Oriental Melittia Hübner (Lepidoptera, Sesiidae), from the collection of Muséum d’histoire naturelle, Genève. Revue Suisse de Zoologie 103:323–338.

Grice, H., A. Freitas, A. Rosa, O. Marini-Filho, F. Dias, N. Mega, M. Casagrande, et al. 2019. Parides hahneli (amended version of 2018 assessment). The IUCN Red List of Threatened Species 2019: e. T16244A145166386.

Hall, D. W. 2014. Hypercompe scribonia (Stoll 1790) (Lepidoptera: Erebidae: Arctiinae). Featured Creatures - Entomology & Nematology, University of Florida.

Hart, N., J. Partridge, I. Cuthill, and A. T. Bennett. 2000. Visual pigments, oil droplets, ocular media and cone photoreceptor distribution in two species of passerine bird: the blue tit (Parus caeruleus L.) and the blackbird (Turdus merula L.). Journal of Comparative Physiology A 186:375–387.

Hart, N. S. 2001. Variations in cone photoreceptor abundance and the visual ecology of birds. Journal of comparative physiology. A, Sensory, neural, and behavioral physiology 187:685–697.

Hernández-Chavarría, F., A. Hernández, and A. Sittenfeld. 2004. The" windows", scales, and bristles of the tropical moth Rothschildia lebeau (Lepidoptera: Saturniidae). Revista de biología tropical 52:919–926.

Holloway, J. D. 1999. The moths of Borneo. Southene Sdn Bhd, London.

Holloway, J. D., G. Kibby, and D. Peggie. 2001. The families of Malesian moths and butterflies. Fauna Malesiana Handbooks (Vol. 3). Brill.

Hoskins, A. n.d. False Methona - Patia orise. Learn about butterflies - the complete guide to the world of butterflies and moths.

Hu, S.-J., X. Zhang, A. M. Cotton, and H. Ye. 2014. Discovery of a third species of Lamproptera Gray, 1832 (Lepidoptera: Papilionidae). Zootaxa 3786:469–482.

Janzen, D. H. 1984. Weather-related color polymorphism of Rothschildia lebeau (Saturniidae). Bulletin of the ESA 30:16–21.

Johnsen, S., and E. A. Widder. 1998. Transparency and visibility of gelatinous zooplankton from the northwestern Atlantic and Gulf of Mexico. The Biological Bulletin 195:337–348.

Maia, R., C. M. Eliason, P. Bitton, S. M. Doucet, and M. D. Shawkey. 2013. pavo: an R package for the analysis, visualization and organization of spectral data. Methods in Ecology and Evolution 4:906–913.

Miller, S. E. 1994. Systematics of the neotropical moth family Dalceridae (Lepidoptera)/mit Abb: Bulletin of the Museum of Comparative Zoology.

Muséum national d’Histoire naturelle. 2003*a*. Gastropacha quercifolia (Linnaeus, 1758). Inventaire National du Patrimoine Naturel.

———. 2003*b*. Thyris fenestrella (Scopoli, 1763). Inventaire National du Patrimoine Naturel.

Negm, H. A. R. 1968. Biology and Ecology of Diaphania Unionalis (Hubner), and Comparative Morphology of D. Unionalis (Hubner), D. Hyalinata (Linnaeus), and D. Nitidalis (Stoll).

N’guessan, K., I. Kébé, and A. Adiko. 2010. Seasonal variations of the population of Eulophonotus myrmeleon Felder (Lepidoptera: Cossidae) in the Sud-Bandama region of Côte d’Ivoire. Journal of applied Biosciences 35:2251–2259.

Nupponen, K. 2015. Interesting records of Ethmiinae from the former USSR, with description of Ethmia ustyurtensis Nupponen, sp. n. from Kazakhstan (Lepidoptera: Gelechioidea, Elachistidae). SHILAP Revista de Lepidopterología 43:125–131.

Orellana, A. 2010. Pyrrhopyginae de Venezuela (Lepidoptera: Hesperioidea: Hesperiidae). Entomotropica 23:177–291.

Orlandin, E., M. Piovesan, F. M. D’Agostini, and E. Carneiro. 2019. Use of microhabitats affects butterfly assemblages in a rural landscape. Papéis Avulsos de Zoologia 59.

Park, B.-S., J.-H. Ko, S.-M. Na, D.-J. Lee, U. Bayarsaikhan, and Y.-S. Bae. 2016. Taxonomic study of the Genus Glyphodes (Lepidoptera: Crambidae) from Laos. 한국자연보호학회지 10:148–154.

Pinna, C., M. Vilbert, S. Borenztajn, W. D. de Marcillac, F. Piron-Prunier, A. Pomerantz, N. Patel, et al. 2020. Convergence in light transmission properties of transparent wing areas in clearwing mimetic butterflies. bioRxiv 2020.06.30.180612.

Pittaway, A. R., and I. J. Kitching. 2000*a*. Cephonodes hylas hylas (Linnaeus, 1771) -- Coffee clearwing; Coffee bee hawkmoth. Sphingidae of the Eastern Palaeartic.

———. 2000*b*. Hemaris radians (Walker, 1856). Sphingidae of the Eastern Palaeartic.

Pomerantz, A., R. Siddique, E. Cash, Y. Kishi, C. Pinna, K. Hammar, D. Gomez, et al. 2021. Developmental, cellular, and biochemical basis of transparency in the glasswing butterfly Greta oto.

Poole, R. W. 1970. Habitat preferences of some species of a Müllerian-mimicry complex in northern Venezuela, and their effects on evolution of mimic-wing pattern. Journal of the New York Entomological Society 121–129.

Project Noah. 2017. Erebid moth - Corematura cf. chrysogastra. Project Noah.

Rab Green, S. B., G. L. Gentry, H. F. Greeney, and L. A. Dyer. 2011. Ecology, natural history, and larval descriptions of Arctiinae (Lepidoptera: Noctuoidea: Erebidae) from a cloud Forest in the eastern Andes of Ecuador. Annals of the Entomological Society of America 104:1135–1148.

Rougeot, P.-C. 1959. Un nouvel Orthogonioptilum du Gabon [Lep. Attacidae]. Bulletin de la Société entomologique de France 64:231–232.

Salazar, J. A., J. Vargas, A. M. Mora, and J. Benavides. 2010. Identificación preliminar de los Rhopalocera que habitan el Centro Experimental Amazónico (CEA) Mocoa-Putumayo y algunas especies aptas para criar en cautiverio (Insecta: Lepidoptera). Bol. Cient. Mus. Hist. Nat. U. de Caldas 14:150–188.

Sastry, C., D. Withington, K. MacDicken, and N. Adams. 1988. Multipurpose tree species for small farm use: proceedings of an international workshop held Nov. 2-5, 1987 in Pattaya, Thailand.

Sbordoni, V., L. Bullini, G. Scarpelli, S. Forestiero, and M. Rampini. 1979. Mimicry in the burnet moth Zygaena ephialtes: population studies and evidence of a Batesian—Müllerian situation. Ecological Entomology 4:83–93.

Seitz, A. 1906. The Macrolepidoptera of the World: A Systematic Description of the Known Macrolepidoptera Edited with the Collaboration of Well-known Specialists (Vol. 6). A. Kernen, Stutgart.

Shapiro, A. M. 1978. Phenotypic and Behavioral Convergence of" Silver-Spotted Skippers"(Lepidoptera: Hesperiidae). Biotropica 10:159–160.

Skinner, B. 2009. Colour identification guide to moths of the British Isles:(Macrolepidoptera). Apollo Books.

Tarmann, G. M. 2005. Zygaenid moths of Australia: a revision of the Australian Zygaenidae (Procridinae: Artonini). Csiro publishing.

Urra, F. 2014. Un nuevo género chileno de Depressariidae (Lepidoptera: Gelechioidea). Boletín del Museo Nacional de Historia Natural, Chile 63:101–110.

Vorobyev, M., and D. Osorio. 1998. Receptor noise as a determinant of colour thresholds. Proceedings of the Royal Society B-Biological Sciences 265:351–8.

Warren, A. D., and N. V. Grishin. 2017. A new species of Oxynetra from Mexico (Hesperiidae, Pyrginae, Pyrrhopygini). ZooKeys 155.

Weller, S. J., R. Simmons, R. Boada, and W. Conner. 2000. Abdominal modifications occurring in wasp mimics of the ctenuchine-euchromiine clade (Lepidoptera: Arctiidae). Annals of the Entomological Society of America 93:920–928.

Willmott, K. R., J. C. R. Willmott, M. Elias, and C. D. Jiggins. 2017. Maintaining mimicry diversity: optimal warning colour patterns differ among microhabitats in Amazonian clearwing butterflies. Page 20170744 *in* (Vol. 284). Presented at the Proc. R. Soc. B, The Royal Society.

Worthy, R., J. M. Gonzalez, and S. D. Rios. 2019. A review of the genus Insigniocastnia JY Miller, 2007 (Lepidoptera: Castniidae) with notes on Castnia amalthaea H. Druce, 1890. Zootaxa 4550:277–288.

Yagi, T., T. Katoh, A. Chichvarkhin, T. Shinkawa, and K. Omoto. 2001. Molecular phylogeny of butterflies Parnassius glacialis and P. stubbendorfii at various localities in East Asia. Genes & genetic systems 76:229–234.

Yazaki, H., M. Kishimura, M. Tsubuki, and F. Hayashi. 2019. Müllerian mimicry between cohabiting final-instar larval Pryeria sinica Moore, 1877 (Lepidoptera: Zygaenidae) and pupal Ivela auripes (Butler, 1877)(Lepidoptera: Lymantriidae). The Pan-Pacific Entomologist 95:83–91.
